# Novel Prion Protein Gene (*PRNP)* Variants in Wild Montana Mule Deer

**DOI:** 10.64898/2026.03.17.711390

**Authors:** Alyssa L. Seerley, Michael T. Rothfuss, Bridget M. Gray, Mylene A. Sebogo, Bethelhem A. Manakelew, June I. Pounder, Bruce E. Bowler, Moses J. Leavens, Andrea L. Grindeland Panter

## Abstract

Chronic Wasting Disease (CWD) is a transmissible spongiform encephalopathy (TSE) of cervids (elk, deer, moose, and reindeer) that is increasing in prevalence and expanding to new geographical areas. TSEs, commonly referred to as prion diseases, are fatal neurodegenerative diseases that occur in a variety of mammals, including humans, and typically exhibit species-specific characteristics. This study reports the sequencing of the prion protein gene *(PRNP)* in retropharyngeal lymph node samples from 358 Montana mule deer (*Odocoileus hemionus*) and the identification of 36 *PRNP* genetic variants, many of which have not been reported previously. Further investigations tracked spatiotemporal characteristics of variants to hunting districts, year of harvest, and CWD status. *PRNP* polymorphisms V12F, D20G, R40Q, and S225F were examined with EmCAST computational predictions to determine the relationship between sequence and structural variations providing further insights into mechanisms affecting CWD outcomes. EmCAST predictions suggest the novel variant V12F phenotype is attributable to functional changes such as altered protein-protein interactions that might be linked to the CWD positive status of the samples. Notably, the analysis of S225F by EmCAST predicted that S225F is a neutral mutation for folded PrP and incompatible with fibril PrP, suggesting a potential structural mechanism for why this previously known variant may provide protection against CWD based on reduced fibril PrP formation. The CWD-positive samples harboring *PRNP* variants were examined with the prion RT-QuIC assay, including the novel variant V12F, which resulted in prion seeding activity.

**Author Summary:** Chronic Wasting Disease (CWD) is a fatal disease of cervids, which include deer, elk, and moose. Since its discovery in 1967, CWD has spread to 36 U.S. states and four Canadian provinces, with prevalences exceeding 20% in select free-ranging populations. With the popularity of hunting big game animals and the role of these species in the ecosystem, concerns have arisen regarding the transmission of disease to humans, as well as how to mitigate long term consequences of disease on animal populations. Given the significant risk of species spillover and the limitations of current management, innovative genetic research is essential. Our study identified novel *PRNP* genetic variants in Montana mule deer, cataloging their regional distribution and CWD status across several hunting seasons. By investigating the impact of these polymorphisms on protein stability and seeding activity, we provide critical insights into the genetic factors that influence disease phenotypes and transmissibility in wild cervid populations.

## 1. Introduction

Transmissible spongiform encephalopathies (TSEs), known as prion diseases, are a class of rare, fatal neurodegenerative diseases that slowly progress over time and have no cure^1^. These diseases can occur in a variety of mammals, including humans, and are typically known for exhibiting species-specific characteristics^1,2^. TSEs can be acquired by three mechanisms: Inherited genetic mutations, spontaneous occurrence with no known etiology, or acquisition of infectious forms of the prion protein^3–6^. Examples of TSEs include Creutzfeldt-Jakob Disease (CJD), Fatal Familial Insomnia (FFI), Gerstmann-Straussler-Scheinker syndrome (GSS), bovine spongiform encephalopathy (BSE), scrapie, feline spongiform encephalopathy, exotic ungulate spongiform encephalopathy, and chronic wasting disease (CWD)^1^.

Prion diseases occur when the normal cell-surface glycoprotein, with an alpha helical conformation (PrP^C^), is converted into a disease-associated conformation containing a high beta-sheet content (PrP^Sc^). This conversion then acts as a template, with PrP^C^ misfolding into PrP^Sc^ aggregates, ultimately leading to neurodegeneration^7–9^. PrP^C^ exists in all mammals where it is expressed throughout the body in organs, tissues, and the central and peripheral nervous systems^9^. Structurally, PrP^C^ has a flexible N-terminal domain, and a C-terminus containing three alpha-helices and two anti-parallel beta-strands flanking the helix^10,11^. While the functional role of the normal form of the prion protein is not well understood, PrP^Sc^ has been extensively studied because of its role in TSEs. An important factor in this process for disease conversion, is the homology between the host’s primary PrP sequence in comparison to the incoming PrP^Sc^. Mismatches between PrP^C^ and PrP^Sc^ sequences may explain prion disease species barriers, however, occasionally prions have been known to “jump” to a different species, including zoonotic transmission^8^. The most notable example of this was the rise of variant CJD (vCJD) in the United Kingdom resulting from oral ingestion of bovine spongiform encephalopathy contaminated meat^7^.

Chronic wasting disease is a prion disease that was first recognized in 1967 in a captive deer in Colorado^2^. This disease affects cervids, which includes, elk, deer, reindeer, and moose, and is the most infective of all prion diseases^2^. CWD is spread among cervids via peripheral shedding of PrP^Sc^ from infected animals through bodily fluids and excrement such as saliva, urine, blood, feces, and additional tissues, including antler velvet before clinical signs of disease appear^1,2,12–16^. Additionally, infectious misfolded prions can persist in the environment long-term independent of a host, bind to vegetation, and potentially infect animals that graze on contaminated foliage^17,18^. The type of vegetation and soil may influence binding properties of the prions, further complicating detection and decontamination in the environment^18–22^. These properties make CWD difficult to manage in wild cervids and may increase difficulty as CWD epidemics progress^23^. Consequently, CWD has been spreading throughout wild populations of cervids and according to the United States Geological Survey (USGS), it is now found in 36 states and 5 Canadian provinces.

Polymorphisms in the prion protein gene (*PRNP)* have been shown to affect host CWD susceptibility, progression of disease, and incubation periods through their encoding of variations in amino acid sequence, informing protein conformation, and thus impacting the conversion of PrP^C^ to the disease-causing form, PrP^Sc^^2,24–29^. Structural elements of PrP differ significantly between genetic variants, with the *PRNP* gene providing templates for prion growth and conformational basis for the propagation of different strains and phenotypes of disease^2,16,30^. Notably, investigations into *PRNP* genetics have historically been useful for designing disease control programs. Researchers discovered that a homozygous genotype for A^136^R^154^R^171^ in sheep and goats conferred resistance to scrapie, an observation that subsequently led to successful eradication of the disease in domestic sheep through breeding programs^2,31–33^. With CWD posing substantial management challenges, researchers have begun to investigate the potential for genetic resistance within susceptible species. Previous studies in mule deer (*Odocoileus hemionus*) have established genetic variations in *PRNP* at amino acid positions 20, 131, 225, 226, and 247^2,8,34,35^, some of which influence susceptibility and progression of disease^8,31^. For example, mule deer have been documented to have a polymorphism at codon position 225, expressing either serine (S) or phenylalanine (F), and this is implicated in variable disease prevalence, altered time course of disease progression, and even variations in disease neuropathology^27,36–38^. A survey of Wyoming and Colorado mule deer show that mule deer with the 225SS *PRNP* variant were 30 times more likely to be CWD positive compared to the animals carrying the 225SF polymorphism^31^. Recently, additional polymorphic *PRNP* variants have been identified in mule deer at codon 20, 96, 98, 116, and 159^29^; however, the downstream PrP structural effects and specific phenotypes are not well understood. As CWD continues to increase in prevalence, the need to investigate emerging genotypes which influence disease progression and species barriers is critical.

In this study we: 1) Identified and cataloged novel and previously known *PRNP* polymorphisms present in Montana’s wild-mule deer population in years 2017, 2018, and 2022; 2) performed a prevalence analysis on the variants using spatiotemporal and CWD data; 3) modeled protein stability and misfolding probability of a subset of the novel variants through EmCAST computational analyses; and 4) confirmed PrP seeding activity of CWD positive protein variants relative to wild-type protein via the use of the Real-Time Quaking-Induced Conversion assay (RT-QuIC).

## 2. Materials and Methods

### ***2.1*** Cervid Samples

Samples of retropharyngeal lymph nodes of wild mule deer (*Odocoileus hemionus*) were provided by Montana Fish, Wildlife & Parks (MT FWP). The 358 samples that were analyzed for this study were collected at MT FWP CWD sampling stations during the 2017, 2018, and 2022 hunting seasons. Samples were screened for CWD via enzyme-linked immunosorbent assay (ELISA) and suspect positives were confirmed positive via immunohistochemistry. All samples used in this study were collected from hunting districts 401, 600, 670, 640, 502, and 555 in Montana with known CWD positive cases (**Figure S1).** The sampling strategy for *PRNP* sequencing included both sexes of deer (2 male:1 female- as male samples were more readily available) equally distributed through each hunting district at all years if samples were available. All CWD positive samples obtained in those years were included in the dataset.

### 2.2 Transgenic Mouse Model Samples

All animal procedures were conducted in compliance with the guidelines outlined in the US National Research Council’s *Guide for the Care and Use of Laboratory Animals*^39^ and the US Public Health Service’s *Policy on Humane Care and Use of Laboratory Animals*. The study protocol was reviewed and approved by the Institutional Animal Care and Use Committee (IACUC) at the McLaughlin Research Institute. Mice were housed in the Animal Resource Center at the Weissman Hood Institute at Touro University, a facility exclusively for mice and accredited by the Association for Assessment and Accreditation of Laboratory Animal Care. This research was approved by the Weissman Hood Institute at Touro University; McLaughlin Research Institutional Animal Care and Use Committee under protocol number 2023-AG-100 and biosafety protocol number 2022-AG-IBC5.

The Tg1536+/+ mouse line, commonly denoted as “cervidized mouse line” was used for controls in the RT-QuIC assay^1,40^. A CWD inoculated group, and normal brain homogenate (NBH) inoculated control group were each included in the study and tissues were harvested at CWD endpoint ages, 9.4 months of age. These mice express cervid PrP at a five-fold higher level than the level of wild-type PrP expression in FVB mice (**Figure S2**) and are maintained on an FVB/NJ background expressing no endogenous mouse PrP. The animals inoculated with CWD were used as a prion-positive control to identify the seeding activity parameters in an overexpressing model. The animals inoculated with NBH were utilized as an additional CWD-negative control. In addition to the CWD and NBH inoculated mice expressing cervid PrP previously mentioned, a control expressing no PrP as a knockout (KO) of the *PRNP* gene was included in the RT-QuIC assay (**Figure S2**). These samples were denoted as “PrP KO mouse brain”. CWD and NBH inoculum were mule deer in origin and confirmed wild-type, lacking *PRNP* polymorphisms.

### 2.3 Sanger Sequencing and Analysis

Frozen lymph node samples were processed for sequencing by standard methods and guidelines as previously reported^41^. 180 microliters (µL) of cell lysis buffer (Tris-Cl pH 8.0, EDTA, NaCl, 0.2% (w/v) sodium dodecyl sulfate) and 20 µL of 10 mg/ml Proteinase K (PK) were added to each well containing a small frozen lymph node sample. Samples were then briefly vortexed and incubated at 50°C for 6 hours. PK digestion was terminated by heating samples at 95°C for 15 minutes. PCR amplification of the targeted *PRNP* region resulted in a 907 bp amplicon. The forward primer was 5-’GCAGTCATTCATTATGCTGCAG-3’ and the reverse primer was 5-’GCAGGTAGATACTCCCTC-3’. Each PCR well contained 13.6 µL of 2x GoTaq cocktail (Promega), 0.405 µL of each primer (20 µM), and 5 µL of diluted DNA (5 µL of lysate in 180 µL of water). The PCR program had 1 cycle of initial denaturation at 95°C for 3 min. Then 35 cycles of denaturation at 94°C for 30 seconds, annealing at 58°C for 30 seconds, and elongation at 72°C for 30 seconds. A final elongation step was done at 75°C for 5 minutes. To check for amplicons, samples were run on a 1.5% agarose gel (0.5X Tris Borate EDTA buffer and Ethidium bromide) at 225V for 35 minutes. Results were visualized and photographed on an ultraviolet transillumination box. Following the PCR amplification, 20µL of the samples were transferred to a 96 well plate for Sanger sequencing by the contract laboratory service provider, Psomagen. Sequence data received from Psomagen was analyzed using Geneious Prime (version 2024.0.2) by aligning and comparing with mule deer sequence GenBank accession number AY228473.1. The sample identification numbers containing single nucleotide polymorphisms (SNPs) which differed from wildtype (WT) *PRNP* were tracked.

### 2.4 Spatiotemporal Distribution of PRNP variants with CWD disease status

The samples expressing the *PRNP* variants were integrated with data provided by MT FWP such as CWD status, hunting district and year of harvest to provide a spatiotemporal assessment. CWD binominal probabilities were calculated for a 95% confidence interval using a calculator that relies on the Clopper-Pearson (exact) method due to small sample sizes (np<5).

### 2.5 Computational Structural Analysis of Mule Deer PrP Variants

EmCAST (empirical Cα stabilization) software models protein sequence-structure relationships in the absence of tertiary interactions using overlapping tetrad fragments^42^. The method is highly sensitive to sequence/structure and can be used to predict short structures, stability changes caused by mutations, and structural distortions caused by mutations^42^. EmCAST uses a sequence-to-structure fragment database to model context-dependent effects in primary structure and to evaluate the different conformations a sequence may encode. Energy calculations are performed for a target protein structure by comparing the population of fragments with matching geometry to the population of fragments with non-matching geometry.

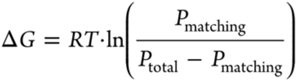

The EmCAST sequence to structure algorithm identifies population peaks for each tetrad (typically 2-4 geometries) and assembles a Cα backbone using every possible combination. Geometric combinations that produce backbone collisions are discarded. The number of generated backbones increases exponentially with sequence length; residues 1-18 of mule deer PrP (MDPrP) produced 98304 backbones. The EmCAST energy function is then applied to all backbone conformations to sort them from lowest to highest energy, as described previously^42^.

To model a larger section of the N-terminal region of MDPrP (Residues 1-44), predictions were performed for three smaller, overlapping fragments (MVKSHIGSWILVLF<u>VAMW</u>, <u>VAMW</u>SDVGLCKKR<u>PKPG</u>, and <u>PKPG</u>GGWNTGGSRYPGQ). Tetrads with few population peaks (VAMW and PKPG) were chosen as the break points. The 50 lowest energy conformations for the three fragments were combined and rescored by EmCAST to generate a predicted lowest energy conformation for residues 1-44. Mutation induced stability changes were calculated using this structure on the EmCAST website (www.emcast.org) with the temperature set to 25°C. The sequence to structure process was repeated for the variants predicted to destabilize MDPrP.

### 2.6 Preparation of Bank Vole Prion Plasmid for RT-QuIC

cDNA encoding Bank vole prion sequence was manufactured by Genscript (Piscataway NJ, USA) and placed in the pET 30a(+) vector (**Table S1**). The cDNA prion sequence was confirmed by Genscript, and the mass of recombinant Bank vole prion protein (residues 23-231, rPrP) was confirmed with electrospray ionization liquid chromatography mass spectrometry (LC-MS) and no heat SDS-PAGE.

### 2.7 Bank Vole Recombinant Prion Expression and Purification for RT-QuIC

Bank vole prion protein is the universal substrate, demonstrating susceptibility to most types of prions, and thus is an ideal substrate for prion seed amplification RT-QuIC assays^43^. Prion RT-QuIC assay was performed as previously described^44^ with modifications as described below. The cDNA encoding Bank vole recombinant prion (rPrP) was expressed from the pET 30a(+) vector by transformation into BL21 *E. coli* competent cells (New England Biolabs, Ipswich MA, USA; DE3, #C2727I). We used our previous protocol to express rPrP from *E. coli*^45^. Sterile 2xYT media (1 L) was inoculated with 10 mL of suspended transformed *E. coli* cells and 1 mL of 50 mg/mL kanamycin and put in a 2.8 L Fernbach flask. Cells were grown in an orbital shaker at 250 rpm and 37°C. One mM final IPTG (GoldBio, #I2481C) was added when OD_600_ was in the range of 0.6 - 0.8. Cells continued to grow for ∼15 hours at 30 °C and 150 rpm. Media was spun for 15 minutes, 7,500 rpm at 4 °C, and cells were frozen at -20 °C. Cells were lysed with 50 mL of 0.5 M NaCl, 0.05 M Tris, pH 8.0 to 5 g cells, placed on ice with 0.5 mg/mL lysozyme, and put on stir plate for 30 minutes, with sonication on ice for 5 minutes (30 seconds on/30 seconds off, 35% AMPL) using a probe sonicator (Sonics VibraCell, Newtown CT, USA). Lysate was spun for 30 minutes at 12,500 rpm and 4 °C, and supernatant was separated from the pellet. The pellet was vigorously disrupted and mixed with ∼50 mL of wash buffer 1 (1% Triton X-114 in 50 mM HEPES pH 8.0), wash buffer 2 (20 mM HEPES 2 M NaCl, pH 8.0), and wash buffer 3 (20 mM HEPES pH 8.0), with a 30 minute, 12,500 rpm (Sorvall RC5B Plus centrifuge, Sorvall SS-34 rotor), 4 °C spin between each wash buffer step. After the spin step with wash buffer 3, the supernatant was discarded, and the pellet was denatured overnight at room temperature in 8 M GuHCl, 20 mM Tris, pH 8.0. The next day, the pellet was visually solubilized and spun at 4 °C, 30 minutes, 12,000 rpm. The supernatant was then added to 20 mL of Ni-NTA resin (ThermoScientific #88222) in a 3 cm (wide) x 30 cm (length) Bio-Rad gravity flow column. After the flow through was collected, the column was washed with 10 column volumes of refolding buffer (100 mM sodium phosphate pH 8.0) and then washed with 10 column volumes of buffer B (100 mM sodium phosphate, 50 mM imidazole, pH 5.8). For the elution step, 10 column volumes of buffer C (100 mM sodium phosphate, 300 mM imidazole, pH 5.8) was added and eluant fractions were collected in ∼14 mL aliquots. Fractions were checked for A_280_ using a Thermo Scientific NanoDrop One spectophotometer. Fractions that contained the highest A_280_ were collected and put in 10,000 MWCO dialysis tubing (ThermoScientific, #68100) overnight at room temperature for ∼18 hours in 10mM sodium phosphate pH 5.8. The next day, dialysate was spun for 30 minutes, 20 °C, and 12,000 rpm. Supernatant was concentrated in a 10,000 MWCO Amicon centrifugal concentrator (Millipore Sigma, Burlington MA, USA) and protein purity and identity was assessed with electrospray ionization LC-MS yielding the expected monomeric mass of 22,914.26 Da (**Figure S3**) and no heat SDS-PAGE stained with Coomassie (**Figure S4**).

### 2.8 Prion RT-QuIC on Mule Deer Lymph Nodes and Brain Homogenates

The deer lymph node tissues were weighed and diluted to 20% w/v using RIPA buffer with protease inhibitor cocktail (Sigma Aldrich #11836170001), then homogenized using a Bead Mill 4 on #5 speed setting for 60 seconds and aliquoted and stored at – 80 °C. Deer and transgenic mouse model brain tissues were homogenized to 10% w/v. The 20% w/v stock homogenates were diluted using 1x PBS to 10% w/v, and then 10-fold dilutions were prepared with 1x PBS/1x N-2 (Gibco, #2389243). The Bank vole rPrP substrate was freshly purified prior to the prion RT-QuIC experiment. After rPrP was dialyzed overnight (∼18 hours), it was spun, concentrated, and filtered with 100 kD Pall centrifugal filters (#OD100C35, LOT #FL6611) at 2,000x*g*, 20 °C, for 10 minutes prior to concentrating to a stock concentration of ∼40 µM or 1.0 mg/mL (ε = 62,005 M^-1^cm^-1^, Thermo Scientific NanoDrop One). The prion RT-QuIC buffer solution was prepared with a 50 mL volumetric flask using 5 mL of 10x phosphate buffer (Boston BioProducts, #BM-220), 45 mL of high-performance chromatography grade water (Sigma Aldrich #270733), 10 µM of EDTA (FisherBiotech #6381-92-6), 300 mM NaCl (Sigma Aldrich, #S9888), 1 mg of SDS (Thermo Scientific, #28365), and 50 mg of potassium phosphate dibasic (FisherBiotech #7758-11-4) to obtain a pH equal to 6.7. For the prion RT-QuIC reaction mix, a stock 5 mM ThT (Thermo Scientific, #211760050) solution in ddH_2_O was added to obtain a 10 µM final ThT in prion RT-QuIC buffer with 0.1 mg/mL (∼4 µM) final concentration of 100 kD filtered rPrP substrate. Ninety-eight microliters of the prion RT-QuIC reaction mix was added to each well in clear bottom 96-well microplates (Thermo-Scientific Nunc 96-well optical bottomed black polystyrene plates w/Lid, catalog #165305) and seeded with 2 µL of 10^-3^ tissue homogenate in each well. Because recombinant substrate can spontaneously convert to amyloid during RT-QuIC reaction conditions, we included rPrP substrate in reaction mix alone as an additional control in the 96 well plate (**Figure S5**). Plates were sealed with tape (Thermo-Scientific, clear polyolefin, non-sterile, catalog #232702) and placed in a FLUOstar Omega reader (BMG Labtech, Germany) at 42 °C, 700 rpm, with 60 seconds of shaking and 60 seconds of resting on a double orbital setting. ThT fluorescence intensity measurements were collected every 40 minutes with 448 nm excitation and 482 nm emission using a gain setting of 1,600. Each sample was replicated in quadruplicate as is standard in this testing method^43^.

### 2.9 Prion RT-QuIC Data Analysis

After the prion RT-QuIC experiment ended, the BMG Omega Mars software .RUC file was exported to Microsoft Excel to process data via SigmaPlot 14.5 or 15.0 (Systat software) for model fitting of prion RT-QuIC kinetic data. We used equation 1, below, as described previously to model the data^45^. In equation 1,

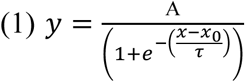

A is ThT fluorescence amplitude, x_0_ is the time to reach midpoint ThT fluorescence, and τ is a fibril time constant where lag phase equals x_0_ – 2τ^45^. Raw kinetic traces were fit to this model using no constraints to estimate x_0,_ τ, and A. The mean ± standard deviation of prion RT-QuIC parameters (ThT fluorescence amplitude midpoint, ThT FA_m_ & lag phase) was then calculated and a Student’s t-test was used to compare mean ThT fluorescence amplitude from CWD cases (n=10) and controls (n=13). The cutoff for RT-QuIC signal was established at 15 hours as no negative controls had positive ThT signal prior to this timepoint. Any sample showing signal after this timepoint was determined negative. A prior ELISA or IHC test performed by the Montana Veterinary Diagnostic Lab on behalf of MT FWP identified these samples as either CWD positive or negative.

For each sample that was ThT positive by RT-QuIC (4/4 wells ThT+), we fit each replicate curve to **Equation 1** to extract ThT fluorescence amplitude, time to midpoint ThT fluorescence, and a fibril time constant, as previously described^45^. A representative figure demonstrating this is shown in **Figure S5.** After fitting to **Equation 1**, we used the prion RT-QuIC parameters (ThT fluorescence amplitude, time to midpoint ThT fluorescence, and the fibril time constant) to calculate lag phase (x_0_-2_τ_) and half ThT fluorescence amplitude (in **Equation 1**, when x = x_0_, y = A/2). With the computed the mean and standard deviation for lag phase and half ThT fluorescence amplitude and then graphed midpoint ThT fluorescence amplitude versus lag phase for each ThT positive PrP^Sc^ biospecimen at 10^-3^ dilution.

## 3. Results

### 3.1 PRNP Sequencing and Identification of Single Nucleotide Variants

The *PRNP* sequences in 358 mule deer samples obtained by MT FWP in the 2017, 2018, and 2022 hunting seasons, were acquired and analyzed resulting in the discovery of 36 polymorphisms. Of these, 25 were non-synonymous *PRNP* genetic variants, many of which, to our knowledge, have not been previously reported in mule deer. These non-synonymous *PRNP* variants resulting in an alternative amino acid at the specified locations were identified in 15.1% of the samples (**Table 1)**. Eleven additional single nucleotide variants identified resulted in synonymous substitutions which resulted in no amino acid translational changes (**Table S2**).

**Table 1:** Non-synonymous *PRNP* polymorphisms (novel and previously reported) identified in Montana mule deer within the study. Previously reported variants were denoted with an asterisk. Variant heterozygote, homozygote frequency and CWD status is reported as a percentage within the polymorphic codon dataset. Sample number reflects the number of samples identified carrying the polymorphic codon out of a total of 358 samples. Binomial probability confidence interval was reported for the CWD positive samples using the Clopper-Pearson (exact) method.

| Polymorphic Codon | Reference Amino Acid | Variant Amino Acid | Reference Sequence | Variant Sequence | Variant Heterozygote Frequency (%) | Variant Homozygote Frequency (%) | CWD Positive (%) | N (out of 358) | Binomial Probability Confidence Interval (95%) |
| --- | --- | --- | --- | --- | --- | --- | --- | --- | --- |
| 2 | Valine (V) | Glycine (G) | GTG | GGG | 100 | 0 | 0 | 1 |  |
| 12 | Valine (V) | Phenylalanine (F) | GTT | TTT | 100 | 0 | 100 | 2 | 0.00068-0.02003 |
| *12 | Valine (V) | Isoleucine (I) | GTT | ATT | 100 | 0 | 100 | 1 | 0.00007-0.01546 |
| 16 | Alanine (A) | Proline (P) | GCC | CCC | 100 | 0 | 100 | 1 | 0.00007-0.01546 |
| 17 | Methionine (M) | Leucine (L) | ATG | TTG | 100 | 0 | 100 | 1 | 0.00007-0.01546 |
| *20 | Aspartic Acid (D) | Glycine (G) | GAC | GGC | 87 | 13 | 14 | 39 | 0.00617-0.03612 |
| 40 | Arginine (R) | Glutamine (Q) | CGA | CAA | 100 | 0 | 0 | 3 |  |
| 52 | Tyrosine (Y) | Serine (S) | TAT | TCT | 100 | 0 | 100 | 1 | 0.00007-0.01546 |
| *96 | Glycine (G) | Serine (S) | GGT | AGT | 0 | 100 | 0 | 1 |  |
| *98 | Threonine (T) | Isoleucine (I) | ACC | ATC | 0 | 100 | 0 | 1 |  |
| 195 | Threonine (T) | Serine (S) | ACC | TCC | 100 | 0 | 0 | 1 |  |
| 204 | Threonine (T) | Asparagine (N) | ACT | AAT | 100 | 0 | 0 | 1 |  |
| 209 | Methionine (M) | Arginine (R) | ATG | AGG | 100 | 0 | 50 | 2 | 0.00007-0.01546 |
| 210 | Glutamic Acid (E) | Aspartic Acid (D) | GAG | GAT | 100 | 0 | 0 | 1 |  |
| 213 | Valine (V) | Glycine (G) | GTG | GGG | 100 | 0 | 100 | 1 | 0.00007-0.01546 |
| 217 | Cysteine (C) | Arginine (R) | TGC | CGC | 100 | 0 | 0 | 1 |  |
| 217 | Cysteine (C) | Serine (S) | TGC | TCC | 100 | 0 | 100 | 1 | 0.00007-0.01546 |
| 221 | Tyrosine (Y) | Serine (S) | TAC | TCC | 100 | 0 | 100 | 1 | 0.00007-0.01546 |
| *225 | Serine (S) | Phenylalanine (F) | TCC | TTC | 100 | 0 | 0 | 2 |  |
| 225 | Serine (S) | Tyrosine (Y) | TCC | TAC | 100 | 0 | 0 | 1 |  |
| 226 | Glutamine (Q) | Arginine (R) | CAG | CGG | 100 | 0 | 100 | 1 | 0.00007-0.01546 |
| 227 | Alanine (A) | Glycine (G) | GCT | GGT | 100 | 0 | 0 | 1 |  |
| 229 | Tyrosine (Y) | Phenylalanine (F) | TAC | TTC | 100 | 0 | 100 | 1 | 0.00007-0.01546 |
| 237 | Leucine (L) | Phenylalanine (F) | CTC | TTC | 100 | 0 | 100 | 1 | 0.00007-0.01546 |
| 238 | Phenylalanine (F) | Leucine (L) | TTC | TTA | 100 | 0 | 0 | 1 |  |

Notably, we frequently discovered multiple polymorphisms within a single deer *PRNP* sequence in our dataset, which is a common occurrence in white-tailed deer^2,24^, yet less frequently reported in mule deer. In fact, all 25 non-synonymous variants were found in a collection of nine deer (**Table S3)**. This representative group is listed with GenBank accession numbers **(Table S3).**

### 3.2 Spatiotemporal Distribution of PRNP Non-synonymous Variants

#### 3.2.1 PRNP polymorphisms identified and spatiotemporal data

The non-synonymous *PRNP* polymorphisms identified in this study were integrated with harvest year and specific Montana hunting region of origin to track how the variant distribution changed over time. Synonymous polymorphisms were not included in the following datasets. In this spatiotemporal analysis, hunting seasons 2017/2018 and 2018/2019 were combined to constitute an “early” timepoint, while hunting season 2022/2023 represents a “late” timepoint. The prevalence of *PRNP* polymorphisms was determined by normalizing the number of samples with variants to the number of samples sequenced in that given hunting district and year. In the early timepoint, both regions 401 and 640 had a similar variant prevalences of 6.5% (4/61) and 6.6% (2/30) respectively. Regions 600 and 670 had variant prevalences of 14% (4/28,) while region 502 had a prevalence of 20% (7/35), respectively. The region with the highest prevalence in the early timepoint was 555 with 33% (10/30) **(Figure 1A).** In the late timepoint, half the regions had similar rates as the early timepoint with region 600, 670, and 502 having 13.6% (3/22), 14.2% (4/28), and 18.5% (5/27) respectively. Region 640 had an increased prevalence of 11% (2/18). The biggest increase was in region 401 that had a variant prevalence of 43.3% (13/30) and the biggest decrease was region 555 with the prevalence of 22.2% (4/18). **(Figure 1B).**

**Figure 1:**
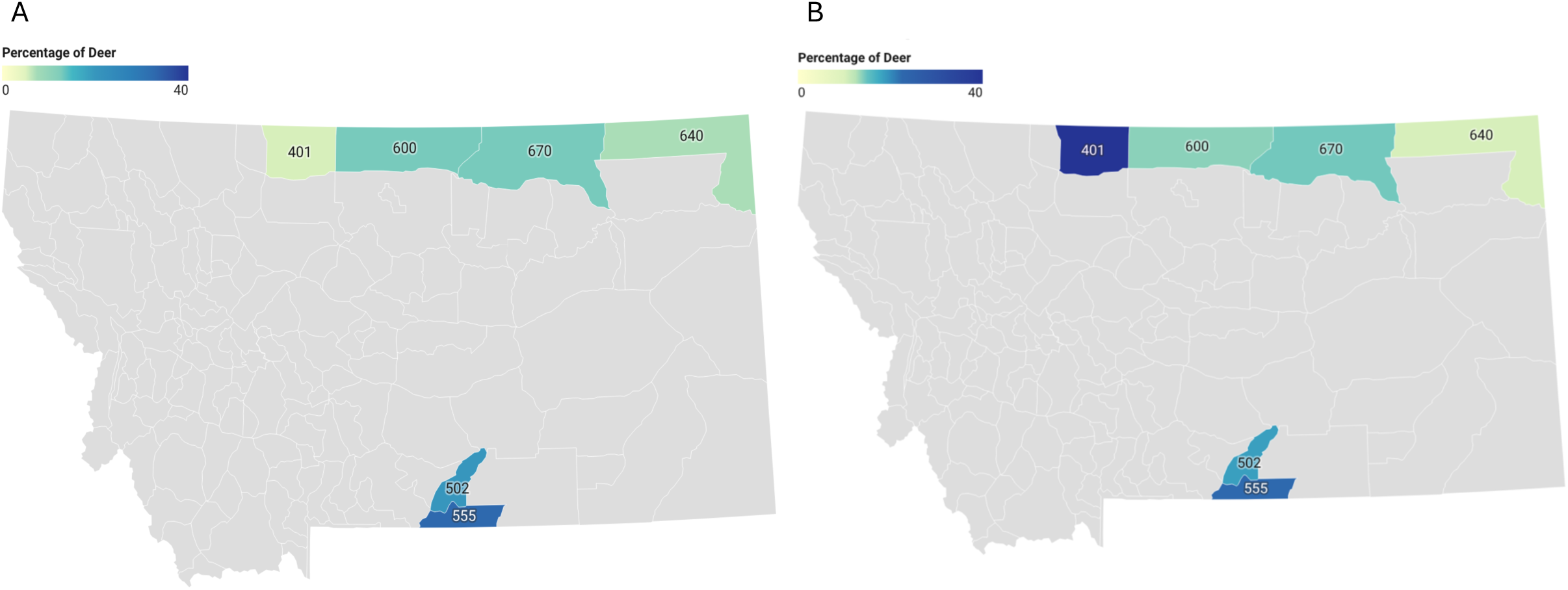
Percentage of mule deer with non-synonymous *PRNP* polymorphisms in relation to hunting district location and time. Prevalence (%) of deer displayed in a heat map carrying a non-synonymous variant in each hunting district from the A) early timepoint hunting seasons (2017/2018 and 2018/2019) and B) late timepoint hunting seasons (2022/2023).

#### 3.2.1 PRNP polymorphisms identified and CWD status

A number of our identified *PRNP* variants were found to be CWD positive, however due to the small sample size, correlations between *PNRP* polymorphism and CWD cannot be determined at this time (**Table 1**). Further investigations were performed, however, on a subset of samples containing both novel and previously reported variants. Our specific criteria for inclusion were: samples containing a variant found in a higher prevalence in the population (three or more) such as codons 20 and 40; samples containing the previously reported variant 225 which has been identified to alter disease phenotypes in the literature^1,31,38^, and polymorphisms which appeared in the central signaling peptide area of the prion protein at codons 2-17.

We examined the overall CWD status of the samples in this condensed dataset which included samples with *PRNP* variants at codons 12, 20, 40, 225, and wild type **(Figure 2 and Table S4)**. The R40Q and the codon 225 variants were not observed in any CWD positive samples; in contrast, all the samples harboring a *PRNP* variant at codon 12 were CWD positive **(Figure 2 and Table S4**). The CWD status for the D20G *PRNP* variants and the wild-type genotype groups were varied, as shown in **Figure 2 and Table S4.**

**Figure 2.**
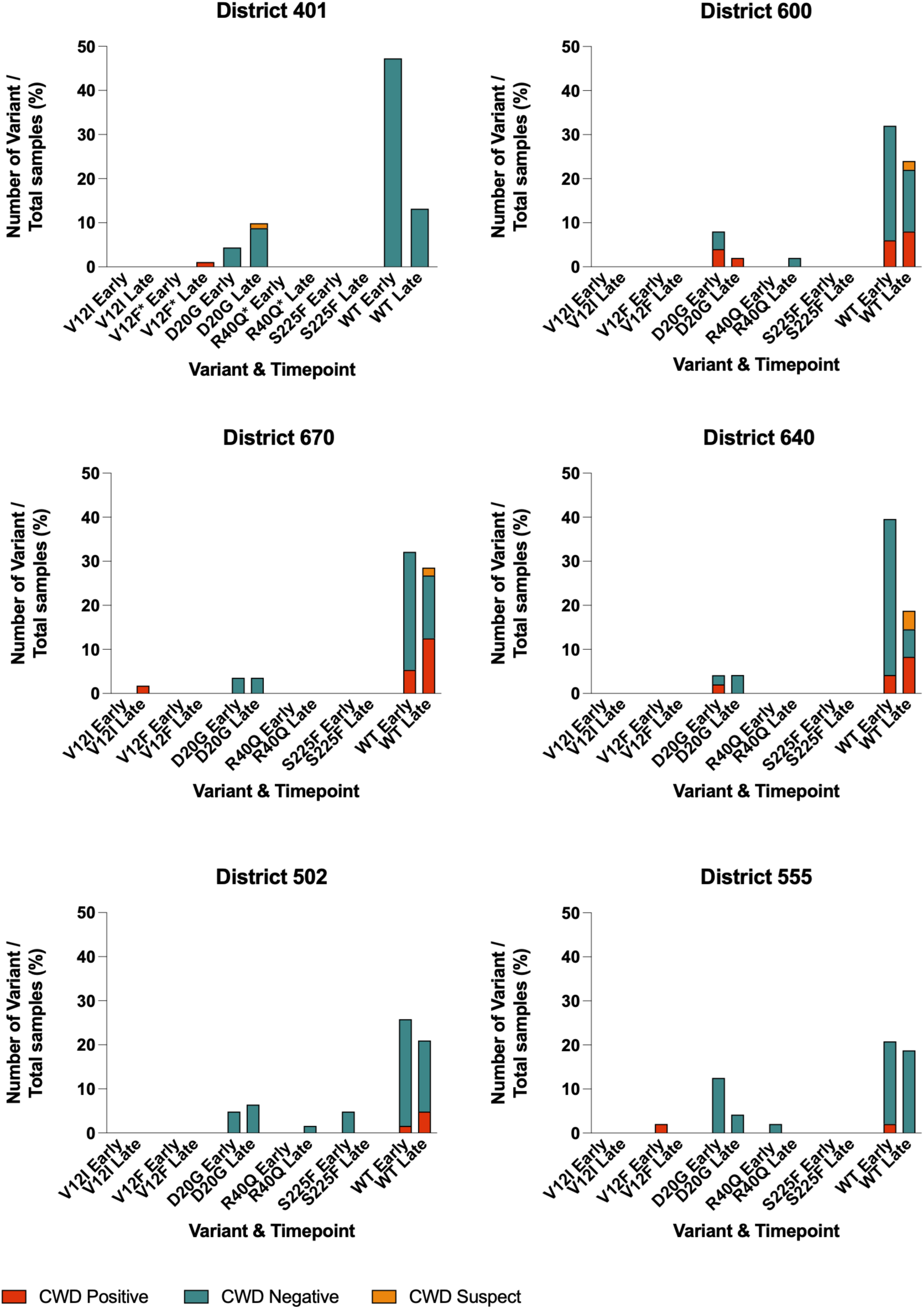
Prevalence of the *PRNP* polymorphisms found to be CWD positive, suspect, and negative within each hunting district and timeframe. *PRNP* non-synonymous variants identified in the hunting districts. Results are shown as percentages of total samples within each hunting district. Red bar graph indicates CWD positive samples, green indicates CWD negative samples, and orange indicates suspect CWD status. This dataset contains a subset of samples of interest; synonymous variants and less common variants were not included.

#### 3.2.1 PRNP polymorphisms, CWD status, and spatiotemporal data

A brief analysis was conducted in our condensed dataset to integrate the *PRNP* polymorphisms discovered, hunting district, timepoint, and CWD status in relation to the total number of samples (358) to better understand the longitudinal trajectories of these variants in wild Montana mule deer.

It was observed that 14 samples were CWD positive in the early timepoint. 10 of these were wild type while the remaining four samples contained one V12F variant and three D20G variants. **(Table S4)**. 135 samples were CWD negative with 113 of them being wild type. Variant S225F appeared two times, R40Q one time, and D20G 19 times **(Table S4)**. The D20G variant appeared in all hunting districts in the early timepoint group and consisted of approximately 3.5%-12.5% of the total samples within each hunting district (**Figure 2**).

The late timepoint group contained 27 total CWD positive samples, 23 of which were wild type. Variant V12F was present once, V12I once, and D20G was present twice **(Table S4)**. One sample with D20G variant was suspected to be CWD positive but not confirmed with further diagnostics. A total of 68 CWD negative samples were within the late group. Forty-nine of those were wild-type, variant R40Q was identified once, while the D20G polymorphism was identified 18 times **(Table S4)**. The D20G variant, was again identified in all hunting districts in the late timepoint ranging from 2%-10% of total samples within each district (**Figure 2)**. Hunting districts 401, 502, and 555 in both timepoints had the greatest number of these variants with 17 out of 25 variants in the early timepoint and 17 out of 24 in the later timepoint **(Figure 2 and Table S4)**.

In addition to examining timing and locality of variants, it was notable to observe that the *PRNP* variants at codon 12 were identified in three different samples that were obtained from three different sample years and three distinct remote areas in Montana, all of which were CWD positive **(Figure 2).** From all three years we sequenced 91 samples in hunting region number 401, and the only CWD positive sample contained our V12F variant. Samples expressing the novel variant, R40Q, were also discovered in different years and hunting regions (**Figure 2 and Table S4)**.

### 3.3 Predictions of Protein Stability Changes Based on PRNP Genetic Variant

We next used computational prediction modeling to probe the stability and structural influence of the single nucleotide alterations which occurred in multiple samples. The variants V12F, D20G, and R40Q occur in the unstructured region of mule deer PrP (MDPrP). EmCAST, a recent tool designed to analyze protein sequence-structure relationships in the absence of tertiary interactions, was used to predict the structural effects of these variants^42^. Sequence-to-structure modeling of the N-terminal region (Residues 1-44) was performed using the wild-type sequence to generate a reference structure (**Figure 3**). A hydrophobic alpha-helix is predicted to span residues 1-27; the remaining residues form a disordered loop. The modeled wild-type structure was used by EmCAST to predict stability changes for the MDPrP variants: V12F (+0.114 kcal/mol), D20G (-0.725 kcal/mol), and R40Q (-0.207 kcal/mol) with positive values indicative of stabilizing impact. Only D20G was predicted to have an impact on stability of the protein specifically. Sequence-to-structure modeling for the D20G sequence predicts D20G to introduce a kink into the N-terminal helix (**Figure 3**). For the S225F variant, EmCAST was run using folded (PDB: 6FNV) and fibril (PDB: 9DMY) structures of MDPrP; EmCAST predicted the S225F variant to be neutral (+0.128 kcal/mol) in the folded structure and incompatible with fibril structure (-Inf kcal/mol).

**Figure 3:**
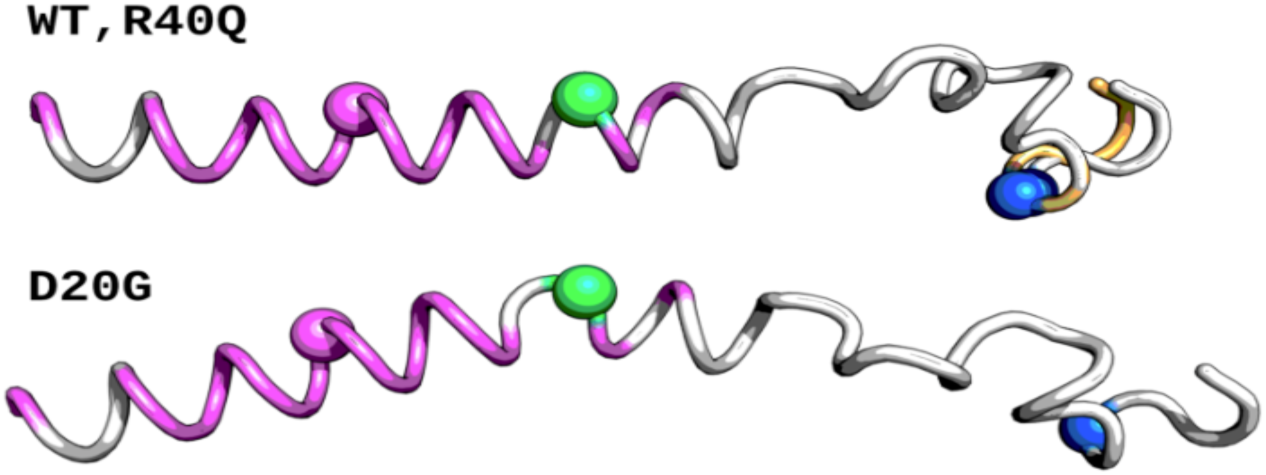
Predicted structures for the N-Terminal region of PrP (Residues 1-44) using WT, R40Q, and D20G sequences in EmCAST. Positions with select variants are rendered as spheres: V12 (magenta), D20 (green), and R40 (blue). Residues 39 and 40-44 are colored orange in the R40Q structure. Hydrophobic residues in the N-terminal helix are colored magenta.

### 3.4 Prion Seeding Activity in Montana Mule Deer Lymph Nodes Expressing Novel CWD-linked PrP Polymorphisms

An exploratory investigation was performed to detect and assess the impacts of a subset of our novel *PRNP* polymorphisms, both with and without CWD, on prion seeding activity. We included a set of novel variant CWD negative samples to both confirm the ELISA results reported for an accurate assessment of the spatiotemporal data, and to investigate the unknown effects the novel variant may have on protein seeding activity. To determine whether PrP^Sc^ was present in a condensed mule deer lymph node dataset expressing the PrP polymorphisms V12F, V12I, D20G, R40Q, and S225F we used the prion RT-QuIC assay to probe lymph node samples along with appropriate controls (**Table 2**)^44^. Samples were chosen based on the presence of our variant of interest; however, we were limited by the samples obtained from the wild specimens. Occasionally, the samples contained more than one polymorphism (**Table 2**). We did not perform RT-QuIC analysis on samples with synonymous single nucleotide polymorphisms, as the amino acid in PrP would be the same as wild-type. We included five controls: 1) brain homogenates from both CWD infected and 2) uninfected brain homogenates of transgenic mice with 4x wild-type cervid *PRNP* (Tg1536+/+)^40^, 3) CWD-positive and 4) CWD-negative mule deer brain homogenates, and 5) *PRNP* knock-out mice. At 10^−3^ dilution, we observed substantial ThT fluorescence in all CWD positive biospecimens (brain or lymph nodes) relative to tissue matched non-CWD controls (**Table 2 and Figure 4**).

**Figure 4:**
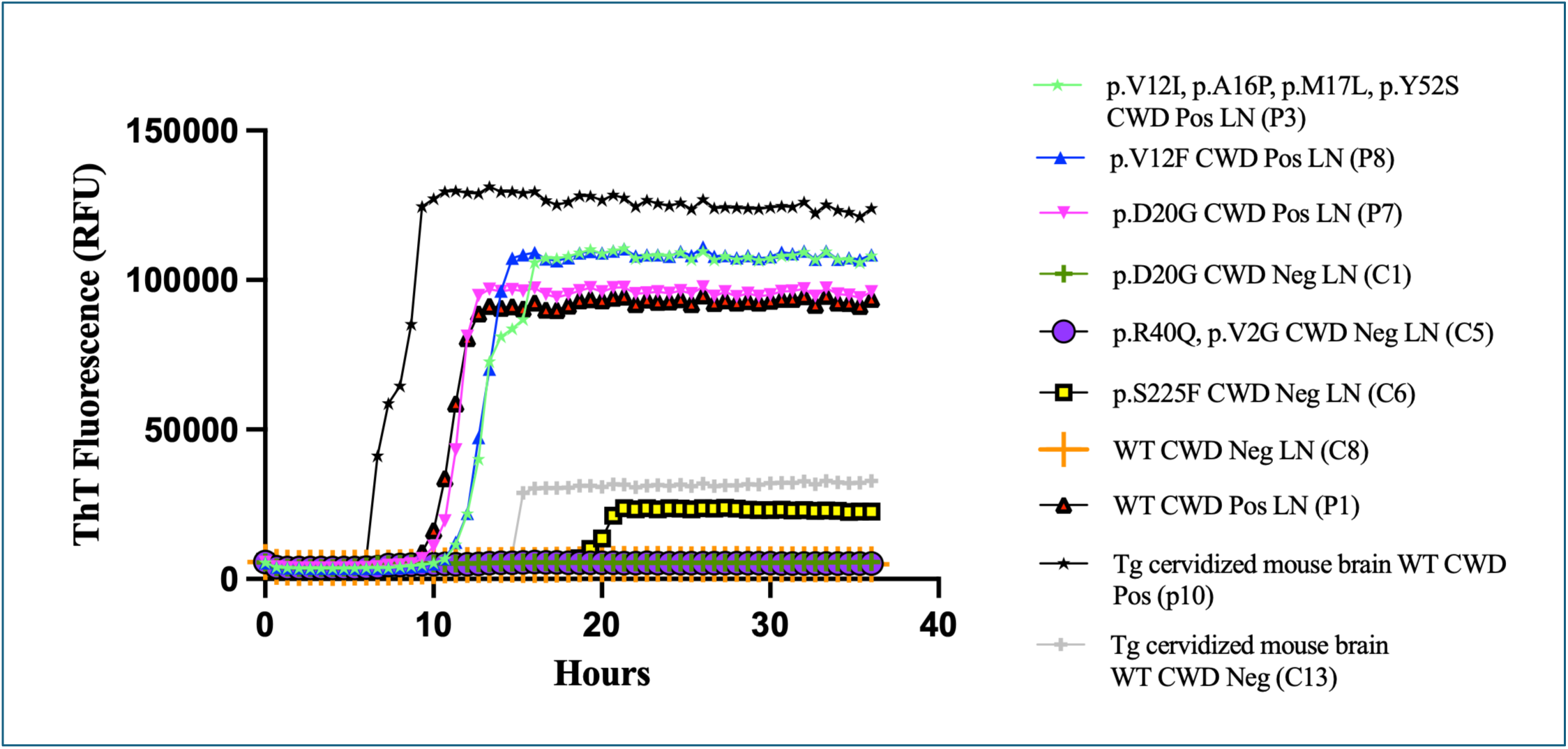
Prion RT-QuIC kinetic data of representative samples by variant group including controls. CWD pos deer lymph nodes expressing novel PrP variants p.V12I, P.A16P, p.M17L, p.Y52S (aqua stars), p.V12F (blue triangles) along with the previously reported variant p.D20G (pink triangles) displaying ThT fluorescent signal. CWD neg novel variant p.R40Q, p.V2G (purple circles), p.D20G CWD neg (green crosses),WT lymph node samples (orange crosses) did not display ThT fluorescent signal. CWD positive transgenic cervidized mouse brain (black stars), was used as a positive control. Tg cervidized mouse brain CWD neg (gray crosses) showed low ThT signal at ∼15 hours; samples after this timepoint were considered negative such as p.S225F (yellow boxes) which displayed low signal after that timepoint.

**Table 2:** RT-QuIC prion seeding activity on mule deer biospecimens harboring *PRNP* polymorphisms. . *Transgenic mouse brain expressing 4x wild-type cervid *PRNP* (Tg1536+/+). Sample Control CWD 10 is a Tg1536+/+ mouse that was inoculated with wild-type CWD positive mule deer brain homogenate and harvested at ∼230dpi. ThT FA_m_ = ThT fluorescence amplitude at the midpoint. CWD status is reported based on previous ELISA results.

| PrP Variant | RT-QuIC Sample Number/Animal ID | Tissue Type | CWD Status | Lag Phase (h) | ThT F<sub>m</sub> (RFU) | Prion RT-QuIC outcome (ThT+/total wells (4)) |
| --- | --- | --- | --- | --- | --- | --- |
| p.D20G | Control 1/4000 (C1) | Lymph Node | Negative | - | - | 0/4 |
| p.D20G | Control 2/295 (C2) | Lymph Node | Negative | - | - | 1/4 |
| p.D20G | Control 3/3489 (C3) | Lymph Node | Negative | - | - | 0/4 |
| p.R40Q, p.D20G | Control 4/47482 (C4) | Lymph Node | Negative | - | - | 0/4 |
| p.R40Q, p.V2G | Control 5/48237 (C5) | Lymph Node | Negative | - | - | 0/4 |
| p.R40Q | Control 10/1358 (C10) | Lymph Node | Negative | 12.4 ± 2 | 40,705 ± 6,901 | 3/4 |
| p.S225F | Control 6/2536 (C6) | Lymph Node | Negative | - | - | 0/4 |
| p.S225F, p.T195S, p.E210D, p.C217R, p.F238L | Control 7/608 (C7) | Lymph Node | Negative | - | - | 0/4 |
| p.WT | Control 8/1006 (C8) | Lymph Node | Negative | - | - | 0/4 |
| p.WT | Control 11/497 (C11) | Lymph Node | Negative | 13.1 ± 2 | 39,708 ± 6,752 | 3/4 |
| PrP Knockout | Control 9/691 (C9) | PrP KO mouse brain | Negative | - | - | 0/4 |
| p.WT | Control 12 deer brain (C9) | Deer brain | Negative | - | - | 0/4 |
| p.WT | Control 13 Tg mouse brain (C13) | Tg cervidized mouse brain* | Negative | - | - | 0/4 |
| p.V12I, p.A16P, p.M17L, p.Y52S | CWD 3 (P3) | Lymph Node | Positive | 11.9 ± 0.7 | 54,300 ± 8,991 | 4/4 |
| p.V12F | CWD 4 (P4) | Lymph Node | Positive | 14.2 ± 2 | 45,200 ± 5,664 | 4/4 |
| p.V12F | CWD 8 (P8) | Lymph Node | Positive | 12.4 ± 1 | 54,100 ± 10,889 | 4/4 |
| p.D20G | CWD 2 (P2) | Lymph Node | Positive | 13.7 ± 0.9 | 55,047 ± 8,288 | 4/4 |
| p.D20G | CWD 6 (P6) | Lymph Node | Positive | 10.5 ± 0.4 | 48,075 ± 10,056 | 4/4 |
| p.D20G | CWD 7 (P7) | Lymph Node | Positive | 11.8 ± 1 | 46,000 ± 5,885 | 4/4 |
| p.WT | CWD 1 (P1) | Lymph Node | Positive | 10.2 ± 0.3 | 43,046 ± 4,005 | 4/4 |
| p.WT | Control CWD 9 (P9) | Deer brain | Positive | 10.4 ± 0.5 | 41,147 ± 3,108 | 4/4 |
| p.WT | Control CWD 10 (P10) | Tg cervidized mouse brain* | Positive | 7.6 ± 1 | 63,300 ± 27,453 | 4/4 |

We observed that the CWD positive transgenic mouse brain expressing 4x deer *PRNP* gave the highest midpoint ThT fluorescence amplitude signal (63,300 ± 27,453) and the shortest lag phase (7.6 ± 1), similar to its kinetic data (Control CWD 10, **Table 2**, **Figure 4 and S6)**. With respect to CWD positive mule deer lymph nodes, the WT, V12F, V12I, and D20G, all displayed significant prion seeding activity via ThT fluorescence amplitude (Student’s t-test, p<0.05) relative to lymph node controls that were ThT negative (**Figure 4 and 5**). The CWD positive mule deer lymph nodes had lag phase values that ranged from 10-14 hours, with midpoint ThT fluorescence amplitude values ranging from 40,000 – 55,000 relative fluorescence units (RFU) (**Table 2 and Figure S6 and S7**). Importantly, the CWD positive mule deer lymph nodes expressing the p.V12F and p.V12I, p.A16P, p.M17L, p.Y52S, demonstrated that these novel polymorphisms had prion seeding activity (CWD samples 4, 8, 3 **Table 2 and Figure 5**).

**Figure 5.**
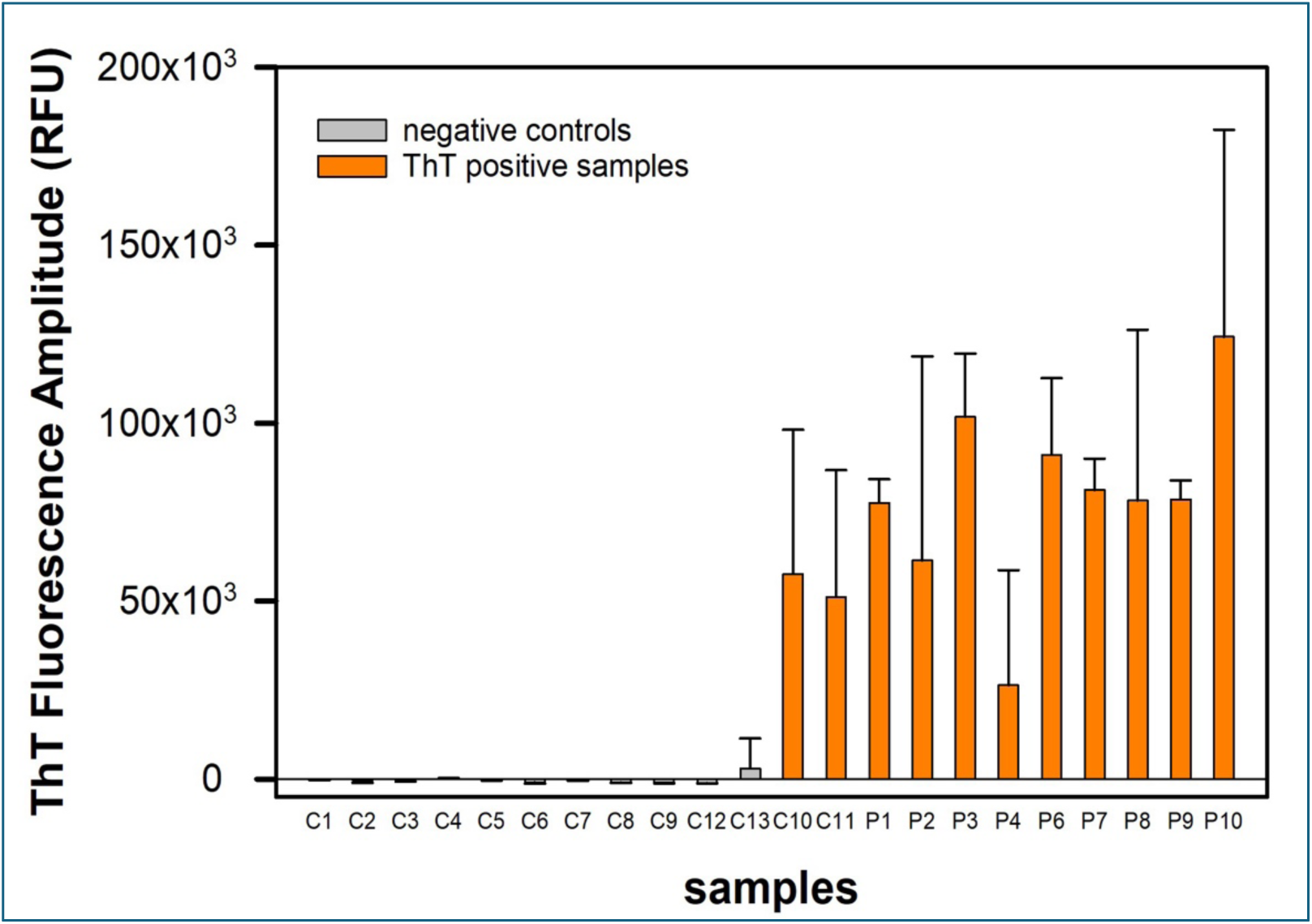
ThT fluorescence amplitude (ThT FA_m_, RFU at 14.6 hours minus initial reading, 0 hours) for samples in study along with controls. Sample identification and abbreviations shown here are denoted in Table 2.

The majority of samples that were determined to be CWD negative via an ELISA, were also ThT negative by prion RT-QuIC, including a *PRNP* knock-out control (Control 9, **Table 2, Figure S8**); however, there were exceptions. Two of the ten mule deer lymph nodes that were determined to be CWD negative by an ELISA, were ThT positive in prion RT-QuIC and had lag phase and midpoint ThT fluorescence amplitude values that clustered with lymph node samples that were CWD positive via ELISA (**Table 2**, **Figure 5, S6 and S9**). One of these two lymph node specimens expressed wild-type PrP (control 11) and the other expressed the PrP variant R40Q (control 10), which to our knowledge, is a novel mutation in mule deer that has been previously reported in sheep^46^. The WT PrP (control 11) had ThT positive signal with a lag phase of 13.1 ± 2 and midpoint ThT fluorescence amplitude of 39,708 ± 6,752 RFU, while the PrP R40Q (control 10) had a lag phase of 12.4 ± 2, and a half ThT fluorescence amplitude of 40,705 ± 6,901 RFU (**Figure S6).**

## 4. Discussion

We report the sequencing of the *PRNP* gene of wild Montana mule deer lymph nodes. We discovered that over half of the mule deer samples possessed at least one of 36 *PRNP* polymorphisms (25 non-synonymous and 11 synonymous) with many samples containing multiple polymorphisms. In this study, we focused on further analysis of a subset of the non-synonymous variants. Nonetheless, we acknowledge that synonymous variants may not only relay information on gene evolution but also may alter protein expression, mRNA splicing and other key properties^47^. Thus, there may be value to further analysis of both the non-synonymous variants not included in our condensed subset and the synonymous variants in future studies.

Prior to the study, we predicted that a positive correlation would occur between the prevalence of CWD and the prevalence of *PRNP* genetic variants. In other words, if CWD prevalence is higher in a region, then selection for *PRNP* variants should be higher if they decrease CWD susceptibility or increase survival time. This was not the case, as regions 600, 670 and 640 had the highest prevalences of CWD, yet did not have significantly higher prevalences of *PRNP* variants (**Figure 2**). Future studies such as this may determine protective genetic variants from CWD driven genetic selection in Montana, however, at the time these samples were procured, detections of disease were relatively new in MT which led to limited sampling. Interestingly, rather than identifying beneficial variants, our data revealed a number of novel *PRNP* variants that were CWD positive, modeled for protein stability, and were confirmed to have PrP seeding activity. Here, we highlight a few polymorphisms investigated in this study.

### 4.1 PRNP polymorphisms within the signaling peptide area of the protein

#### 4.1.1 V12F

The novel genetic variant V12F is intriguing due to the location of the substitution in the center of the signaling peptide, and the discovery that both deer in which it was identified were CWD positive **(Figure 6)**. Two different *PRNP* V12F samples were discovered, and they were obtained from different sample years and distinct remote areas in Montana. Intriguingly, our newly identified V12F variant was the only CWD positive sample out of 91 samples in hunting district 401. The observation of this variant in different years and distinct regions of Montana suggests that it was not transmitted from one generation to the next. An explanation for this could be either that it has arisen multiple times spontaneously, a product of the movement of the wild mule deer, or that it is present in geographical areas that link the distinct regions but was not detected. Because this variant is rare in our dataset, and because we only sampled in several geographically distinct areas, we cannot reliably comment on the distribution or frequency of this variant across the rest of Montana or its association with CWD.

**Figure 6.**
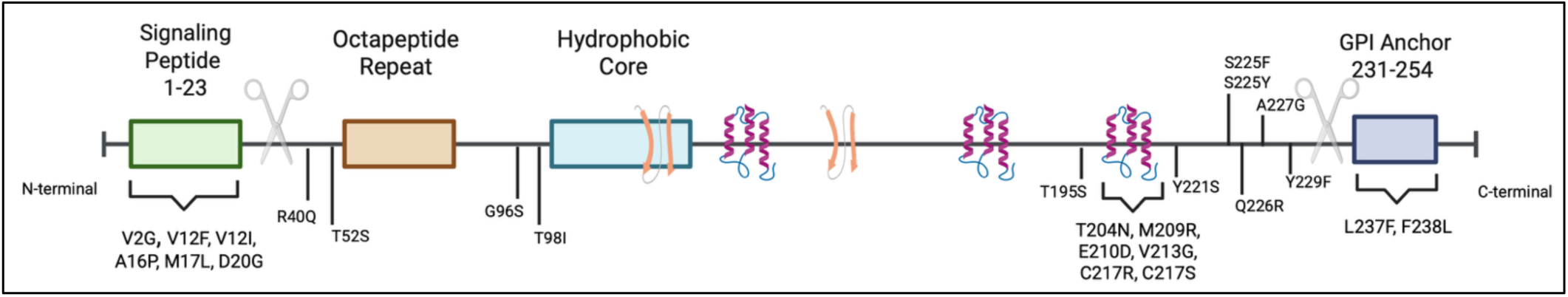
Linear schematic representation of the cellular prion protein (PrP) highlighting key regions, structures, and where all 25 non-synonymous variants are found.

We hypothesized that these single non-synonymous nucleotide substitutions may alter the PrP folding characteristics, and therefore, disease outcomes. Investigations into PrP indicates that amino acids 12, 15, and 16 reside in the N-terminal region of the immature protein prior to cleavage during posttranslational modifications which is a region not thought to have an impact on protein misfolding in prionopathies **(Figure 6)**^48^. Immature PrP has an N-terminal signal peptide at amino acids 1-23 that codes for entry into the endoplasmic reticulum of the cell^49–51^. The N-terminal region of PrP is natively unstructured and its function is in endocytic trafficking of PrP; this region also includes a sequence that is essential for toxicity of PrP^52^. Alterations to the protein structure in the N-terminus may affect intracellular trafficking, may induce specific biochemical changes resulting in PrP misfolding, or may lower the threshold for PrP corruption from another CWD positive animal. Deer with the V12F *PRNP* variant were heterozygous, expressing phenylalanine in one allele and the wild-type valine in the other allele. Valine is known to contain a small hydrophobic side chain as opposed to phenylalanine with a large hydrophobic aromatic side chain which likely disrupt interactions with a binding partner important for proper trafficking^53^. Thus, this variant could disrupt structure causing steric clashes, cause enhanced interactions with other hydrophobic residues that promote conversion, and/or alter protein stability, all leading to misfolding^54^.

When we turned to modeling structural changes, the analysis predicted that the amino acid change from valine to phenylalanine at amino acid 12 was slightly stabilizing; however, this individual amino acid change is not the only factor to consider in protein folding/misfolding predisposition. EmCAST modeled p.V12F to be a structurally neutral mutation in a hydrophobic alpha-helix suggesting that the V12F phenotype might be attributable to functional changes such as altered protein-protein interactions that might be linked to the CWD positive status of all V12F-containing samples. Additionally, the increased hydrophobic nature of phenylalanine enhances the likelihood of conversion to a misfolded protein, as it is an important driver for endoplasmic signal peptide recognition and any changes to this may have an impact on PrP processing^55,56^.

When we directly tested whether there was disease-associated prion seeding activity in the lymph nodes from Montana mule deer expressing the PrP variant V12F, we found prion seeding activity. The detection of prion seeding activity in mule deer lymph nodes expressing this V12F polymorphism via cell-free seed amplification (i.e., RT-QuIC) analysis has not been previously reported. We also recognize that the sample size of this variant is low (N=2) and that further investigations are needed to determine CWD susceptibility in relationship to this variant.

#### 4.1.1 V12I

Notably, another *PRNP* genetic variant was discovered in a single sample at codon 12 which included the four polymorphisms V12I, A16P, M17L, Y52S and was CWD positive by ELISA and RT-QuIC. Three of these prion protein variants in this sample, V12I, A16P, M17L, occur within the center of the signaling peptide of PrP **(Figure 6)**. All three samples with polymorphisms in this area of the protein were CWD positive, providing further evidence that these variants might disrupt proper trafficking resulting in failure or perturbations of normal PrP signal peptide cleavage. Prion seeding activity was confirmed in the RT-QuIC assay.

Due to the small sample size, it is unknown at this time if the *PRNP* variants found in the signaling peptide acquired bona-fide CWD or if a spontaneously misfolding event occurred. Future investigations are critical in determining the relationship of polymorphisms in this area of the prion protein and CWD.

#### 4.1.1 D20G

This variant is also located within the signaling peptide of PrP and was the most commonly found non-synonymous polymorphism, appearing in 12% of the samples **(Figure 6)**. The D20G polymorphism was present in all hunting districts from each season (**Figure 2**). Previous mule deer reports of the D20G variant are conflicting as to whether CWD susceptibility is altered due to the single nucleotide polymorphism at this location^2,31,57,58^. In our study, the disease prevalence within the D20G group at 16.2% was similar to wild type at 20.3% suggesting this is a neutral substitution pertaining to CWD. In other words, substitutions in the signal peptide outside of the central area may be trafficked properly and the signal peptide is cleaved off, therefore, is processed similarly as the WT PrP.

Deer with at least one copy of the allele of the common genotype expressing aspartic acid were less likely to test positive in studies conducted in Canada and Nebraska, as opposed to deer sampled in Wyoming and Colorado where they report the substitution at codon 20 appeared to have no definitive correlation with CWD status^2,57^. The copy number of the D20G polymorphisms was not discussed in these studies. Notably, our investigation identified five homozygote animals out of a total of 39 carrying this variant. We observed that copy number did not appear to influence CWD disease status as both positive and negative samples were discovered within the homozygous samples.

To better understand the D20G variant effect on CWD status^2,31,57,58^, we modelled the polymorphism using EmCAST. The modeling predicted that this variant introduces a kink at the C-terminus of the helix while leaving the hydrophobic region of the helix mostly unperturbed (**Figure 3**). This variant is not known to affect properties such as protein-protein interactions in this region, which may explain inconsistencies between previous studies.

Our RT-QuIC data confirmed prion seeding activity was present in all of our CWD infected samples expressing the PrP D20G variant and was not observed in the CWD negative samples harboring the p.D20G.

### 4.2 R40Q

Another novel *PRNP* variant discovered in this study is at amino acid 40 that substitutes an arginine for glutamine and was identified in three samples. The *PRNP* R40Q variant has been reported in Awassi sheep, but to date, it has not been reported in mule deer populations^46^. Due to the small sample size containing this variant, strong conclusions cannot be made. However, it is interesting to note that only one sample with this variant was identified in 2017 whereas two were identified in 2022. The different samples with this variant were obtained from different hunting regions, two in the south (from years 2017 and 2022) and one in the north (from 2022). While not definitive, this implies that the variant was not genetically inherited between the animals. Following the prevalence of this variant will provide insight into its relationship with CWD disease susceptibility.

EmCAST computational predictions placed R40 in a disordered loop with only minor perturbations predicted for the R40Q variant. This polymorphism occurs in a disordered region of the protein, making it more difficult to predict structural impact, and producing conflicting results regarding whether the variant renders the protein more or less stable. The R40Q mutation removes a charged residue. It is noteworthy that charge patterning can affect the properties of disordered protein regions in ways that may cause the disordered region to be more susceptible to aggregation or conversion^59,60^. With all samples (*n*=3) being initially identified as CWD negative by ELISA, further investigations on impacts of the change in PrP structure in relation to CWD are warranted.

The R40Q variant samples were particularly noteworthy pertaining to the prion seeding activity data. All of the samples containing this variant were reported to be CWD negative by ELISA. However, we observed lag phases and midpoint ThT fluorescence amplitude values that were similar to those of CWD positive samples expressing the p.V12F, p.D20G, and WT PrP proteins (**Table 2**, **Figure 5, Figure S9**). The prion seeding activity in these deer lymph nodes harboring this polymorphism could suggest that this variant might play a role in CWD susceptibility or pathology. Additionally, the RT-QuIC assay is known to be very sensitive^44,61–63^, and therefore may have detected CWD at a low level or early stage of disease, yet the ELISA failed to detect CWD. Lastly, as a universal substrate, the bank vole substrate is known to be highly sensitive and may have led to a spontaneous ThT signal as has been reported previously^64^. This possibility is unlikely, as additional negative controls in the RT-QuIC runs remained negative. Future studies will quantify the seeding dose from tissues expressing these novel PrP variants, and whether there is similar prion infectivity relative to wild-type protein.

### 4.3 S225F

The *S225F* variant is well documented and has been reported to alter disease phenotypes in mule deer^2,31,38^. Specifically, in previous studies, animals expressing the S225F polymorphism have been shown to have reduced probability of acquiring CWD and an altered phenotype of disease, suggesting increased stability of the PrP folding. This variant was present in our population; however, the prevalence was extremely low. In our Montana deer sample set, only two deer were identified with this p.S225F polymorphism, which is in line with previous studies that report this variant as rare^29^, especially when the disease is new to an area. Both deer, one female and one male were found in the 502 district of Montana during the 2017-2018 season. While we lack data for stronger conclusions at this time, the observation that out of 358 samples, only two deer were identified with this polymorphism leads us to speculate whether this variant is less likely to arise independently than other variants that were identified in multiple years and completely different districts. Given that this variant has been shown to provide CWD resistant properties, further investigations into its characteristics could provide helpful insight into how this variant might influence population level CWD dynamics, further informing CWD management. Notably, previous investigations into the mechanisms of PrP misfolding due to sequence and structural variations in the S225F variant are limited^65^. However, analysis of S225F by EmCAST predicted that S225F is a neutral mutation for folded PrP and incompatible with fibril PrP, suggesting a potential structural mechanism for why this variant may provide protection against CWD^2,31,38^ based on reduced fibril PrP formation.

### 4.4 WT

Overall, the seeding activity of the *PRNP* wild-type samples reflected the CWD status reported by ELISA, however, one of our ELISA reported CWD negative samples did have ThT signal via the RT-QuIC assay. All replicates (4/4) of this sample resulted in positive ThT signal, therefore, we believe this sample is CWD positive and the ELISA test result is not in agreement with the RT-QuIC assay. As RT-QuIC is reported to be a more sensitive test than ELISA in cervid retropharyngeal lymph nodes^63^, it may have detected the presence of CWD, while the ELISA reported a false negative. Sample control 8, which was also a WT CWD negative sample, did not contain seeding activity.

## 5. Conclusion

CWD is an infectious prion disease currently limited to cervids (elk, deer, moose, and reindeer) with no current evidence of transmissibility to humans^64,66–68^. However, with CWD spreading and becoming an epidemic among deer in the United States and around the world^69^, scientists remain concerned about the future potential for CWD to spill over into other species, including humans^69^. The host expression of *PRNP* is required for PrP^C^ to PrP^Sc^ conversion and sequence variations may greatly alter phenotypes, affect disease transmission, and influence species barriers^2,7,8,27,65^. The effect of these reported polymorphisms in mule deer *PRNP* is not greatly understood and may lead to new prion strains^70^ with unknown disease outcomes. Evidence of the potential of CWD spillover to other species in laboratory settings^71–73^, demonstrates one of many important reasons to identify and investigate downstream effects of *PRNP* genetic variants harbored in wildlife. Our study contributes valuable information on *PRNP* variants identified in wild mule deer, the timing and geographical location of the variants and the effects on prion protein folding and seeding activity. With the identification of 35 total polymorphisms, these studies contribute to the framework of knowledge regarding CWD nationally and provides valuable information regarding CWD management decisions in the state of Montana.

## Acknowledgments

Research reported in this publication was supported by the National Institute of General Medical Sciences of the National Institutes of Health under Award Number P20GM152335, the Weissman Hood Institute; McLaughlin Research Institute institutional start-up funding, and the Jorgensen Foundation. EmCAST analysis was supported by the National Institute of General Medical Sciences of the National Institutes of Health under Award Number R01GM148610 (B.E.B.)

The authors would like to acknowledge Montana Fish, Wildlife & Parks for providing the tissues and associated demographic and disease data. We thank Emily Almberg, Bevin McCormick, and Austin Wieseler for discussions regarding the manuscript content and Sam Treece and Austin Wieseler for assistance with sample organization and acquisition.

We would like to thank Renee Reijo Pera and the COBRE administrative core for thoughtful discussions regarding this manuscript. We would thank Teresa Gunn and the McLaughlin Research Institute -Gene Editing and Mouse Models Assessment (GEMMA) Core Facility within the Center for Integrated Biomedical and Rural Health Research, RRID:SCR_027045, 1P20GM152335 for sequencing analysis training and conversations about our investigations on the relationship between *PRNP* genetics and PrP protein misfolding. Additionally, thanks to Macie Frans for RT-QuIC figure creation.

We also thank Brent Race for scientific mentoring and discussions regarding CWD and *PRNP* polymorphisms.

This manuscript is dedicated to Tim Wylder, our good friend who believed in this study. Our time together spent discussing this project and other topics will be profoundly missed.

