## Supplementary Material for "Novel Prion Protein Gene (*PRNP)* Variants in Wild Montana Mule Deer"

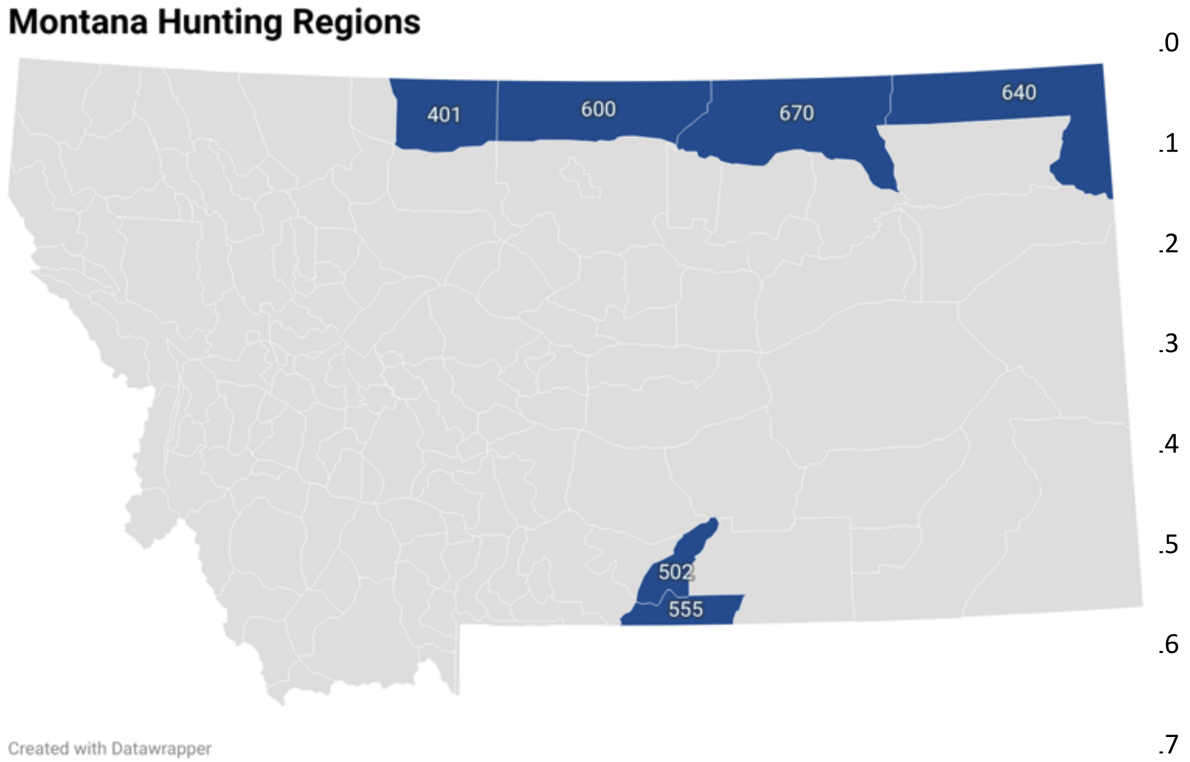

**Figure S1**. Montana hunting regions from which cervid samples in this dataset were collected.

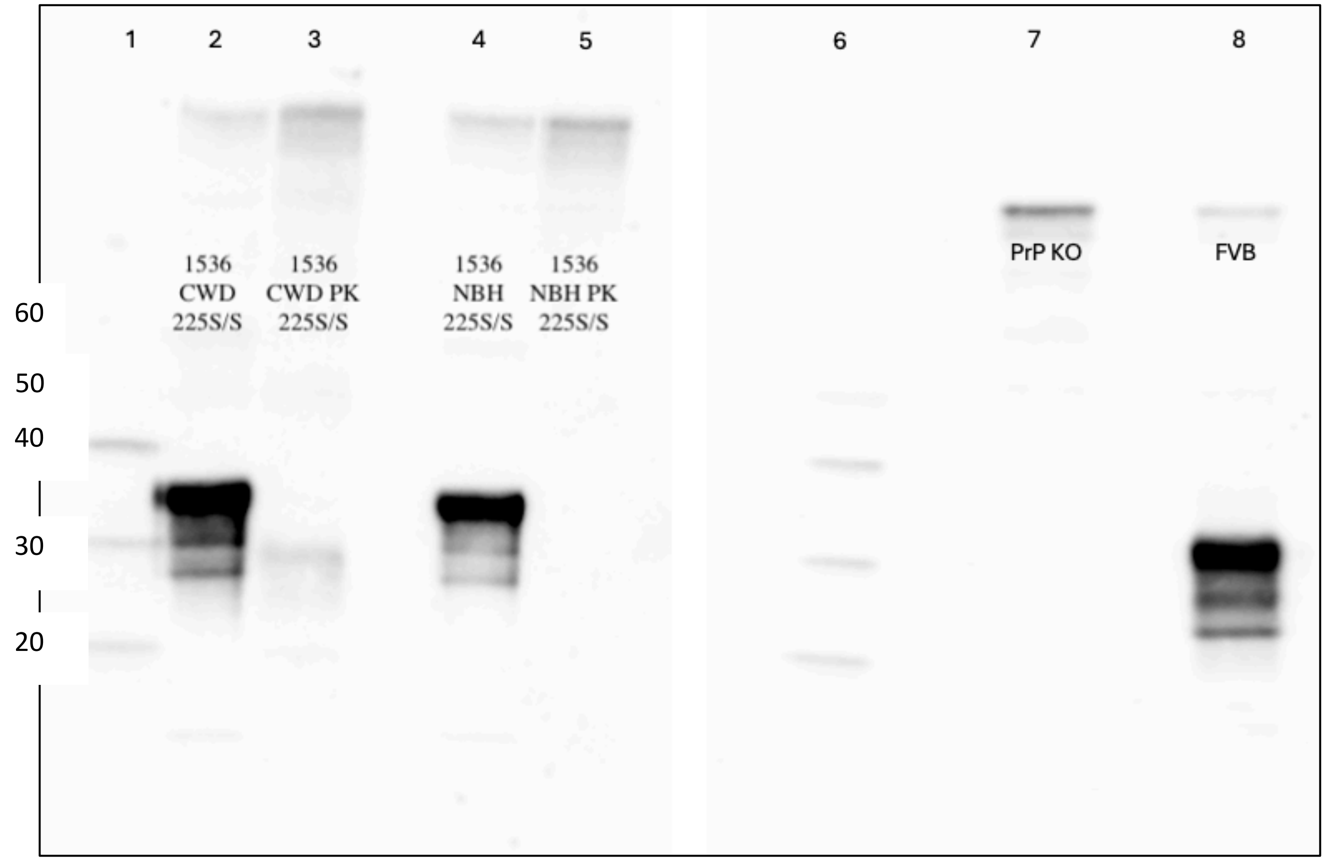

**Figure S2: Transgenic mouse validation of PrP expression and infection using Western blot probed with 6D11**. Lanes 1 and 6 are MagicMark XP standard. Lane 2 is CWD inoculated Cer-PrP-225SS (Tg1536+/+). Lane 3 is CWD Cer-PrP-225SS (Tg1536+/+) with PK treatment to display PK resistant prions. Lane 4 is normal brain homogenate (NBH) inoculated Cer-PrP-225SS (Tg1536+/+) control. Lane 5 is NBH Cer-PrP-225SS (Tg1536+/+) with PK treatment displaying the absence of PK resistant prions. Lane 7 is PrP KO and lane 8 is FVB/NJ expressing endogenous mouse PrP only (no cervid PrP).

5’-
CATATGAAAAAGAGGCCAAAGCCGGGTGGTTGGAACACCGGTGGCTCTCGCTACCCGGGCCAAG
GCTCCCCGGGTGGCAACCGCTACCCACCGCAGGGCGGCGGCACGTGGGGTCAACCGCACGGTGGT
GGCTGGGGTCAACCGCACGGCGGTGGGTGGGGTCAACCTCACGGCGGCGGTTGGGGGCAACCGC
ACGGCGGTGGTTGGGGCCAGGGTGGTGGCACCCATAACCAGTGGAATAAGCCGAGCAAACCGAA
GACCAATATGAAGCATGTTGCAGGTGCTGCGGCGGCGGGTGCGGTGGTGGGTGGCCTGGGCGGTT
ATATGTTGGGCAGCGCAATGAGCCGTCCGATGATCCACTTTGGCAACGACTGGGAAGATCGTTAC
TACCGCGAGAACATGAATCGTTACCCGAATCAGGTGTATTATCGTCCGGTTGACCAGTATAACAAC
CAAAACAACTTCGTGCATGATTGTGTTAATATCACCATTAAACAACATACCGTAACTACGACGACC
AAAGGTGAGAACTTCACCGAAACCGACGTTAAAATGATGGAAAGAGTTGTCGAGCAGATGTGCGT
GACCCAGTACCAGAAAGAGAGCCAGGCCTATTACGAAGGTCGTTCGTAAAAGCTT -3’

**Amino acid sequence**

**N-terminus**-MKKRPKPGGWNTGGSRYPGQGSPGGNRYPPQGGGTWGQPHGGGWGQPHGG
GWGQPHGGGWGQPHGGGWGQGGGTHNQWNKPSKPKTNMKHVAGAAAAGAV
VGGLGGYMLGSAMSRPMIHFGNDWEDRYYRENMNRYPNQVYYRPVDQYNN
QNNFVHDCVNITIKQHTVTTTTKGENFTETDVKMMERVVEQMCVTQYQKE
SQAYYEGRS-**C terminus**

**Table S1. Bank vole cDNA and amino acid sequence information.**

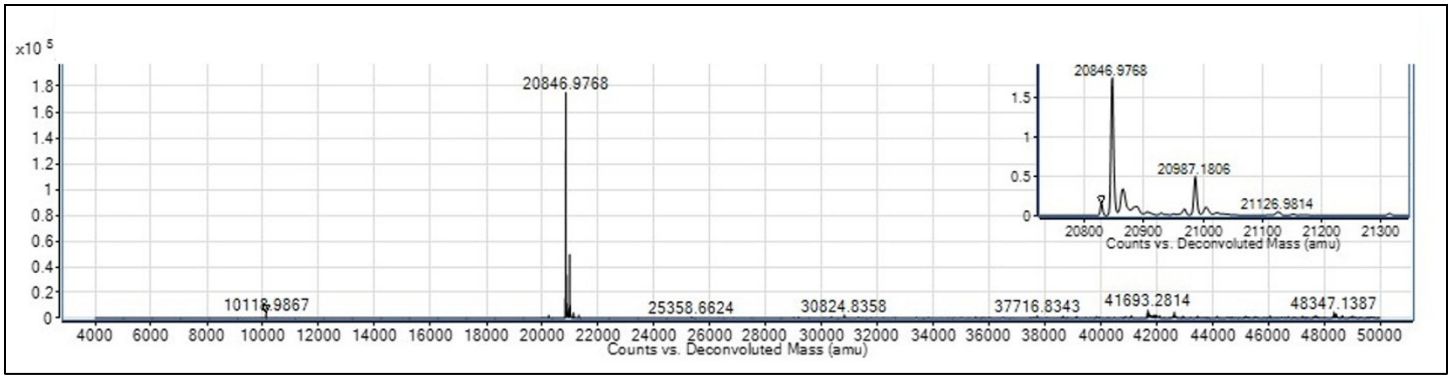

**Figure S3. Electrospray ionization LC-MS deconvoluted mass spectrum of purified Bank vole prion protein in 25 mM Tris pH 8.0.** Expected theoretical mass is 22,914.26 atomic mass unit (from ExPASy Peptide Mass tool (http://web.expasy.org/peptide_mass/) using average mass, with no cutting and $[M+H]^+$ options).

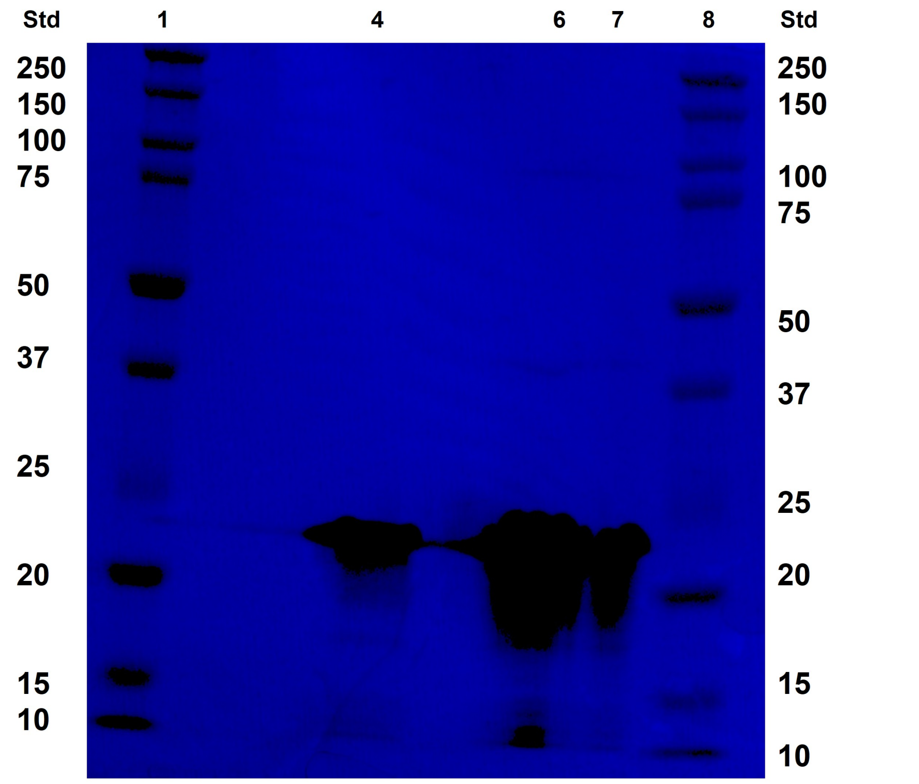

**Figure S4. No heat SDS-PAGE of recombinant Bank vole prion protein (residues 23-231) stained with Coomassie blue.** Lanes 1 and 8 are 10 µL of Biorad precision plus dual color standard, lane 4 is 100 kD filtered Bank vole rPrP at 15 µM used for RT-QuIC experiment, lanes 6 and 7 are unfiltered Bank vole rPrP at 40 µM and 15 µM.

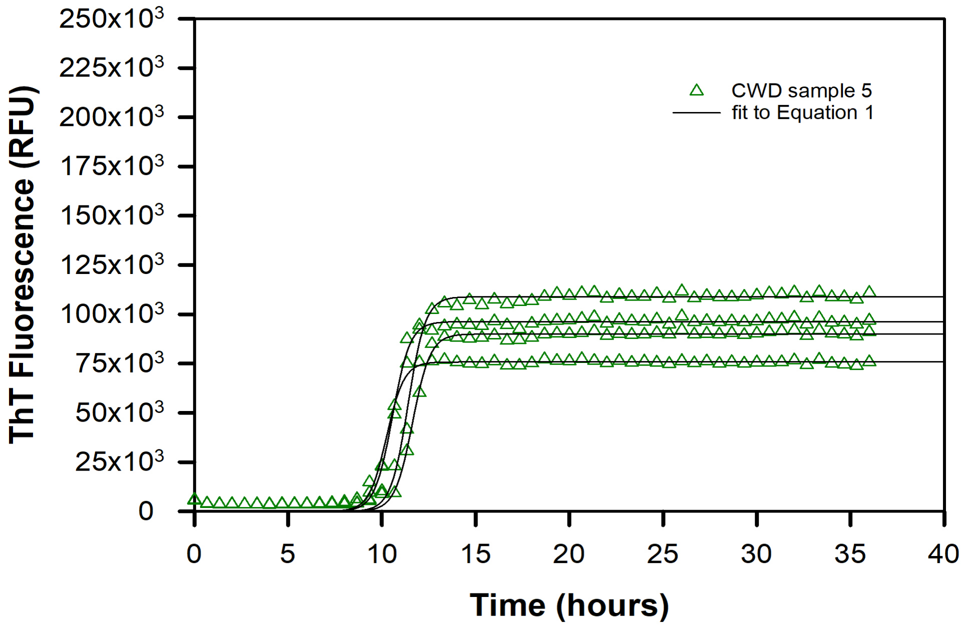

**Figure S5. Representative figure of CWD positive data fitted to sigmoidal model (Eq 1 in main text) to estimate lag phase and ThT amplitude.**

| Polymorphic Codon | Reference Amino Acid | Variant Amino Acid | Reference Sequence | Variant Sequence | Variant Heterozygote Frequency (%) | Variant Homozygote Frequency (%) | CWD Positive (%) | Sample Numbers (out of 358) |
|---|---|---|---|---|---|---|---|---|
| 25 | Lysine (K) | Lysine (K) | AAG | AAA | 100 | 0 | 0 | 1 |
| 108 | Proline (P) | Proline (P) | CCA | CCG | 0 | 100 | 0 | 1 |
| 131 | Tyrosine (Y) | Tyrosine (Y) | TAC | TAT | 93 | 7 | 26 | 96 |
| 139 | Arginine (R) | Arginine (R) | AGG | AGA | 0 | 100 | 0 | 4 |
| 156 | Asparagine (N) | Asparagine (N) | AAT | AAC | 0 | 100 | 25 | 6 |
| 185 | Isoleucine (I) | Isoleucine (I) | ATC | ATT | 0 | 100 | 0 | 1 |
| 202 | Threonine (T) | Threonine (T) | ACT | ACC | 17 | 83 | 25 | 6 |
| 237 | Leucine (L) | Leucine (L) | CTC | CTG | 100 | 0 | 0 | 1 |
| 238 | Phenylalanine (F) | Phenylalanine (F) | TTC | TTT | 83 | 17 | 18 | 12 |
| 247 | Isoleucine (I) | Isoleucine (I) | ATC | ATT | 93 | 7 | 31 | 14 |
| 253 | Leucine (L) | Leucine (L) | CTC | CTT | 100 | 0 | 0 | 1 |

**Table S2: Synonymous *PRNP* polymorphisms identified in the dataset. Heterozygote, homozygote frequency and CWD percentages are within the polymorphic codon group.**

| Polymorphic Variant | Genbank ID (Accession number) |
|---|---|
| V2G R40Q | PZ139486 |
| V12I A16P M17P Y52S | PZ123357 |
| V12F | PZ123356 |
| D20G R40Q | PZ139487 |
| D20G T204N M209R S225Y A227G | PZ139637 |
| G96S | PZ139635 |
| T98I | PZ139636 |
| T195S E210D C217R S225F F238L | PZ139489 |
| M209R V213G C217S Y221S Q226R Y229S L237F | PZ139488 |

**Table S3:** Representative list of *PRNP* polymorphisms discovered within study, some of which carried multiple polymorphisms within the gene.

| | | AA p.V12I | | AA p.V12F* | | AA p.D20G | | AA p.R40Q* | | AA p.S225F | | Wild-Type | |
|---|---|---|---|---|---|---|---|---|---|---|---|---|---|
| Year | Hunting Region | CWD + | CWD - | CWD + | CWD - | CWD + | CWD - | CWD + | CWD - | CWD + | CWD - | CWD + | CWD - |
| 2017 | 401 | 0 | 0 | 0 | 0 | 0 | 2 | 0 | 0 | 0 | | 0 | 21 |
| | 502 | 0 | 0 | 0 | 0 | 0 | 3 | 0 | 0 | 0 | 2 | 1 | 15 |
| | 555 | 0 | 0 | 1 | 0 | 0 | 6 | 0 | 1 | 0 | | 1 | 9 |
| | 600 | N/A | | N/A | | N/A | | N/A | | N/A | | N/A | |
| | 640 | N/A | | N/A | | N/A | | N/A | | N/A | | N/A | |
| | 670 | N/A | | N/A | | N/A | | N/A | | N/A | | N/A | |
| 2018 | 401 | 0 | 0 | 0 | 0 | 0 | 2 | 0 | 0 | 0 | 0 | 0 | 23 |
| | 502 | N/A | | N/A | | N/A | | N/A | | N/A | | N/A | |
| | 555 | N/A | | N/A | | N/A | | N/A | | N/A | | N/A | |
| | 600 | 0 | 0 | 0 | 0 | 2 | 3 | 0 | 0 | 0 | 0 | 3 | 13 |
| | 640 | 0 | 0 | 0 | 0 | 1 | 1 | 0 | 0 | 0 | 0 | 2 | 17 |
| | 670 | 0 | 0 | 0 | 0 | 0 | 2 | 0 | 0 | 0 | 0 | 3 | 15 |
| 2022 | 401 | 0 | 0 | 1 | 0 | 1 | 8 | 0 | 0 | 0 | 0 | 0 | 12 |
| | 502 | 0 | 0 | 0 | 0 | 0 | 4 | 0 | 1 | 0 | 0 | 3 | 10 |
| | 555 | 0 | 0 | 0 | 0 | 0 | 2 | 0 | 0 | 0 | 0 | 0 | 9 |
| | 600 | 0 | 0 | 0 | 0 | 1 | 0 | 0 | 1 | 0 | 0 | 6 | 7 |
| | 640 | 0 | 0 | 0 | 0 | 0 | 2 | 0 | 0 | 0 | 0 | 6 | 3 |
| | 670 | 1 | 0 | 0 | 0 | 1 | 2 | 0 | 0 | 0 | 0 | 8 | 8 |

**Table S4: Mule deer non-synonymous *PRNP* polymorphisms (condensed dataset) integrated with CWD status in Montana hunting regions and year of harvest.** *Novel mule deer *PRNP* polymorphisms are denoted with an asterisk. Red numbers indicate that at least one sample was suspected of CWD. Blue numbers indicate that at least one sample had multiple variants at another position which may affect total sample number. Samples found with synonymous polymorphisms not included.

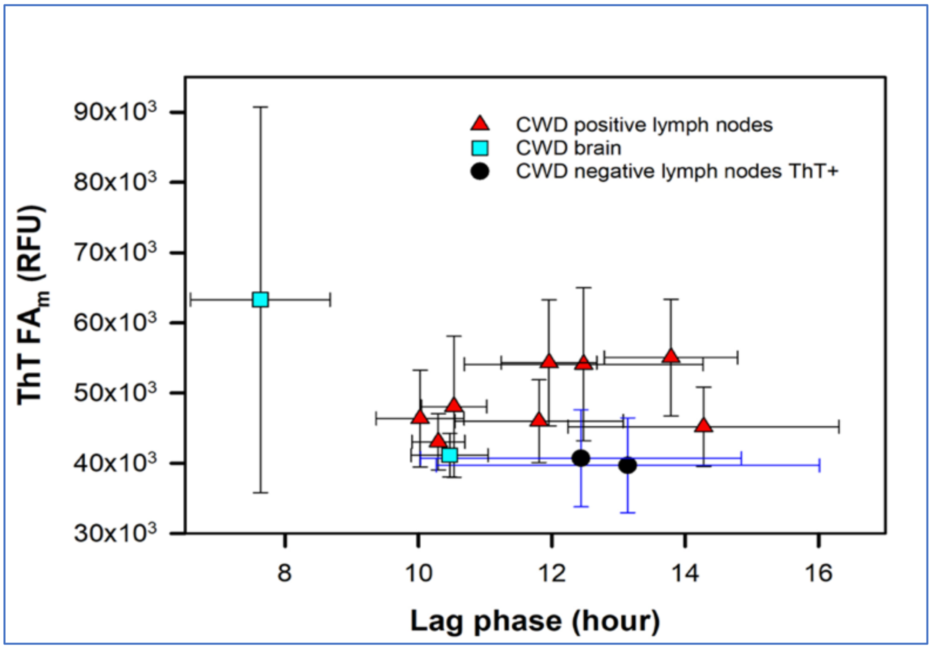

**Figure S6: Lag phase (hours) versus half ThT fluorescence amplitude at the midpoint (ThT FA$_m$, RFU) for all ThT positive biospecimens via prion RT-QuIC.**

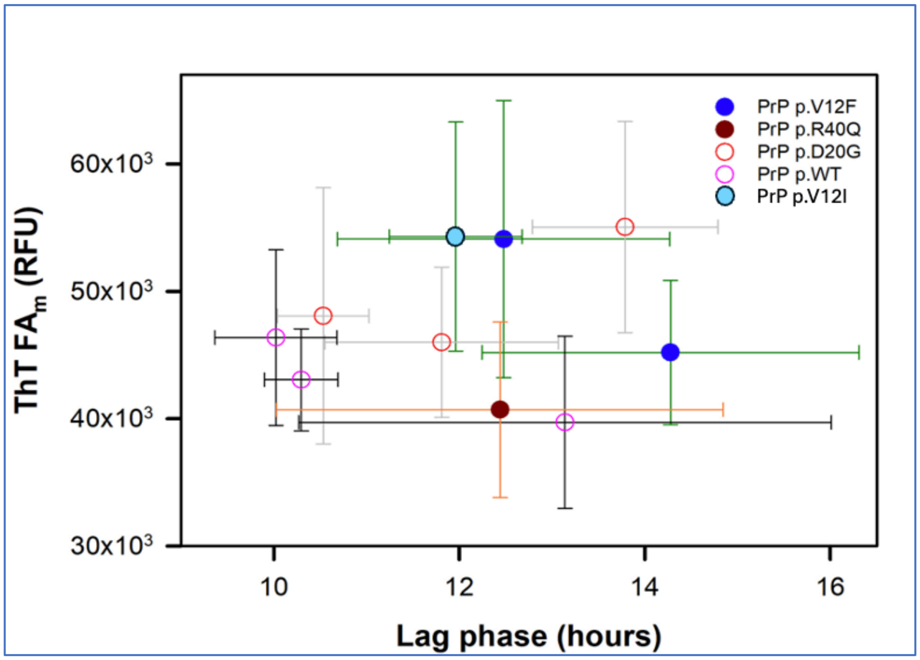

**Figure S7. PrP variant and wild type sample comparisons of ThT fluorescence via lag phase (hours) and midpoint ThT fluorescence amplitude (ThT FA$_m$, RFU).**

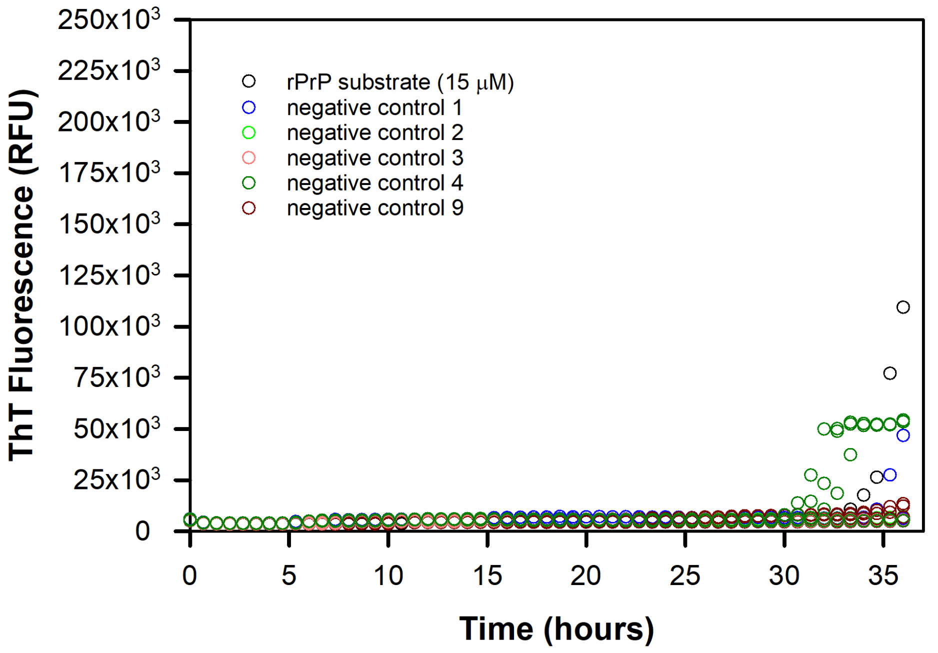

**Figure S8. Prion RT-QuIC assay displaying lack of ThT signal in CWD negative samples and controls**. CWD negative samples 1, 2, 3, 4 (deer lymph node) and 9 (mouse brain *PRNP* knock-out) at $10^{-3}$ dilution.

91

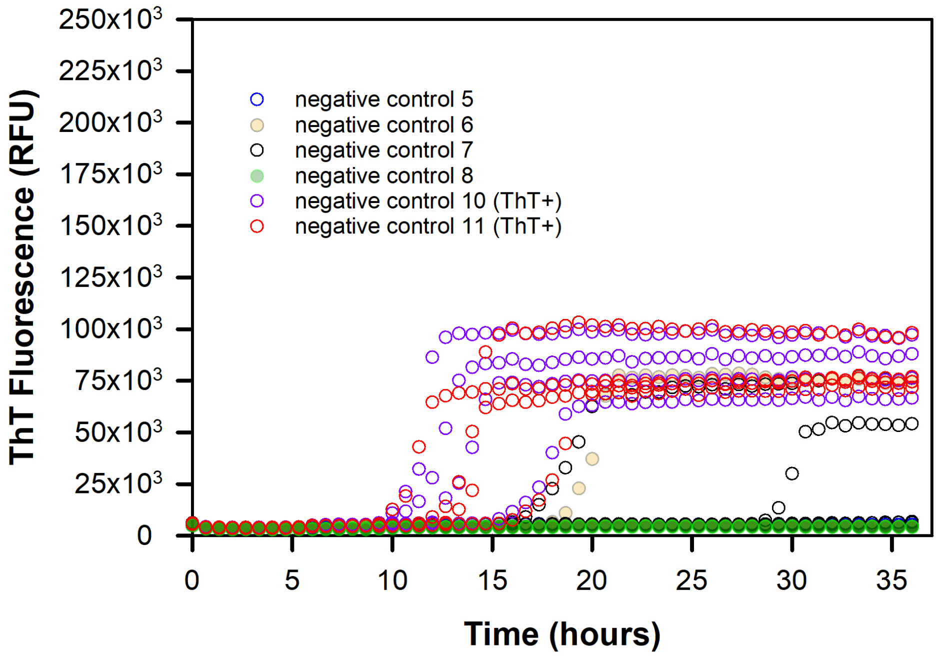

**Figure S9. Prion RT-QuIC assay on samples 5, 6, 7, 8, 10, 11 (lymph node) all reported CWD negative by ELISA at $10^{-3}$ dilution**. Replicates of samples 10 and 11 display ThT positive signal before 14.6 hours.
